# A non-lemniscal thalamic interface connecting alerting sensory cues to internal states in mice

**DOI:** 10.1101/2022.06.19.496703

**Authors:** Yiwei Wang, Ling You, KaMun Tan, Meijie Li, Jingshan Zou, Wenxin Hu, Tianyu Li, Ruizhi Yuan, Fenghua Xie, Fengyuan Xin, Miaomiao Liu, Yixiao Gao, Congping Shang, Zhiwei You, Xiaorong Gao, Wei Xiong, Peng Cao, Minmin Luo, Feng Chen, Bo Hong, Kexin Yuan

## Abstract

Alterations in internal states, such as elevated arousal level and increased anxiety or fear, triggered by alerting environmental cues are required for behavioral state transitions promoting survival. However, the specific brain region that plays an interfacing role between alerting stimuli and internal states remains to be identified. Here, we report that the medial sector of the auditory thalamus (ATm), which consists of a group of non-lemniscal thalamic nuclei, can fulfill this function. VGluT2-expressing ATm (ATm^VGluT2+^) neurons receive direct and strong inputs from both visual and auditory midbrain regions, and project to multiple downstream structures critically involved in brain state regulation. Their activity was correlated with, and indispensable for, both blue light- and sound-induced NREM sleep-to-Wake transition, and their arousing effects were mainly mediated by, but not limited to, the temporal association cortices. ATm^VGluT2+^ neuron activation in awake behaving mice induced pupil dilation and behavioral responses suggestive of anxiety. Blocking the neurotransmitter release of ATm^VGluT2+^ neurons receiving auditory inputs selectively abolished loud noise-triggered escape behavior but not locomotion. Thus, the ATm is an interface in mouse brain that can transform alerting environmental cues into internal arousal and emotional state alterations that promote survival.

## Introduction

Proper brain states are required for adaptive expression of behaviors in ever-changing environment. Brain state at any given time can be determined as a point in a high-dimensional space ^1^, and arousal is one of the dimensions that has undergone extensive studies ^2, 3, 4^. This is not only because arousal is relatively easy to be observed externally, but also because it powerfully modulates perception, cognition, and behavior ^1^. Through decades of studies, it is clear now that arousal level is strongly regulated by intrinsic factors such as circadian rhythms ^5^, energy homeostasis ^6^ and sleep homeostasis ^7^. Besides, changes in arousal level are also commonly observed when one is exposed to behaviorally meaningful environmental cues, especially the alerting ones ^8^, ^9^. Furthermore, since dimensions in the brain state space are often interdependent or correlated, the variations in arousal level are often concomitant with those in other dimensions such as the valence of emotion ^10^. How altering sensory cues trigger coordinated shifts in the two important dimensions, arousal and emotion, of brain state is still poorly understood.

The thalamus is considered as a critical hub region in the brain ^11^. The sensory division of the thalamus is particularly relevant with the two questions raised above, because it receives direct sensory inputs from the periphery and relays sensory information to a variety of cortical and subcortical targets. Within the sensory thalamus, subnuclei are generally divided into the following two types: lemniscal (first-order) and non-lemniscal (higher-order) thalamic nuclei ^12, 13, 14^. Neurons in the lemniscal thalamic nuclei, such as the ventral division of the medial geniculate body (MGBv) and the dorsal lateral geniculate nucleus (dLGN), predominantly respond to sensory stimulus of a single modality. As the last stop before stimulus feature-related information reaches the cortex, the lemniscal sensory thalamus has been thought to play a central role in gating conscious perception during sleep ^15^. Interestingly, a seminal work conducted within the auditory cortex of marmoset monkeys demonstrated that sound-driven responses averaged across neuronal population did not significantly decrease when the animal fell asleep ^16^, consistent with the findings made by PET imaging in human subjects ^17^. Since the primary sensory cortices are predominantly activated by their corresponding lemniscal thalamic nuclei, these results suggest that the latter may not contribute to sensory evoked arousal. Echoing with this assumption, optogenetic stimulation of the ventrobasal complex (VB), the primary somatosensory thalamus, in mice failed to induce sleep-to-wake transition in two recently published independent studies ^18, 19^, although microarousals were reported in another study using similar stimulation approach ^20^.

Compared with neurons in the lemniscal thalamus, those in the non-lemniscal division exhibit remarkably different characteristics. For example, they receive inputs from cortical and subcortical sensory regions of multiple modalities ^21, 22, 23^, and many of them are preferentially activated by novel stimuli ^24^, suggesting that they are suitable for detecting sudden environmental changes. The medial sector of the auditory thalamus (ATm), which consists of the posterior intralaminar thalamic nuclei, peripeduncular nucleus, medial division of the MGB and suprageniculate nucleus, has long been defined as a part of the non-lemniscal auditory thalamus ^25, 26^. Most of the neurons in the ATm express VGluT2 (ATm^VGluT2+^). These neurons receive direct and strong inputs from the superior and inferior colliculus ^21^, which are visual and auditory midbrain, respectively. In turn, they make dense projections to both the temporal association cortex and ectorhinal cortex (TeA/ECT), which are extensively connected with cortical and subcortical brain regions associated with arousal ^27^, and to subcortical regions mostly known for emotional and behavioral state modulation, including but not limited to the ventromedial hypothalamus (VMH), tail of the striatum (TS), bed nucleus of the stria terminalis, lateral amygdala, basomedial amygdala and medial amygdala ^21^. Based on this unique input-output architecture of ATm^VGluT2+^ neurons, we hypothesize that the ATm serves as an interface between external sensory world and internal brain states, and that it may play a pivotal role in mediating sensory evoked behavioral state transitions.

Here, we monitored the activity of ATm^VGluT2+^ neurons by calcium imaging via fiber photometry ^28^ and arousal level via electroencephalography (EEG) and electromyography (EMG) recordings in sleeping mice or pupillometry in awake mice, in conjunction with optogenetic manipulation at circuit level for reversible control of neural activity ^29, 30^ and a constellation of behavioral assays. We found that, in contrast to lemniscal thalamic neurons that showed state-independent population responses, ATm^VGluT2+^ neurons strongly responded to sound or blue light only when mice were awakened from NREM sleep or awake. Photoactivating ATm^VGluT2+^ neurons induced arousal both during NREM sleep and wakefulness, manifested by NREM sleep-to-wake transition and pupil dilation, respectively. Photoinhibition assay revealed that ATm^VGluT2+^ neuronal activity or their signaling to the pyramidal neurons located in deep TeA/ECT was indispensable for sensory evoked arousal in both modalities, whereas the contribution of their signaling to tested subcortical targets showed modality preference. Besides arousal, ATm^VGluT2+^ neuronal excitation in awake mice elicited anxiety- and fear-like behaviors, suggesting an induction of negatively valenced emotional state. Finally, blocking the neurotransmitter release of ATm^VGluT2+^ neurons receiving auditory inputs selectively abolishes white noise-induced escape behavior without affecting locomotion. These findings indicate that the ATm is an interface transforming alerting sensory signals into arousal associated with defensive emotional state.

## Results

### Sensory responsiveness of ATm^VGluT2+^ but not lemniscal thalamic neuronal population correlates with sleep-wake states

If ATm^VGluT2+^ neurons indeed play a critical role in sensory evoked sleep-to-wake transition, their responses to sensory stimuli during sleep ought to be distinct between trials that mice are and are not awakened from sleep. To test this hypothesis, we unilaterally injected Cre-dependent AAV encoding Ca^2+^ sensor GCaMP6m (AAV2/9-hSyn-DIO-GCaMP6m) into the ATm of VGluT2-IRES-Cre mice to selectively express GCaMP6m in VGluT2+ neurons (Fig. 1a), followed by the implantation of accessories for EEG and EMG recordings and of optical fiber targeting the ATm for photometry recordings in freely-moving mice (Fig. 1b).

**Fig. 1.**
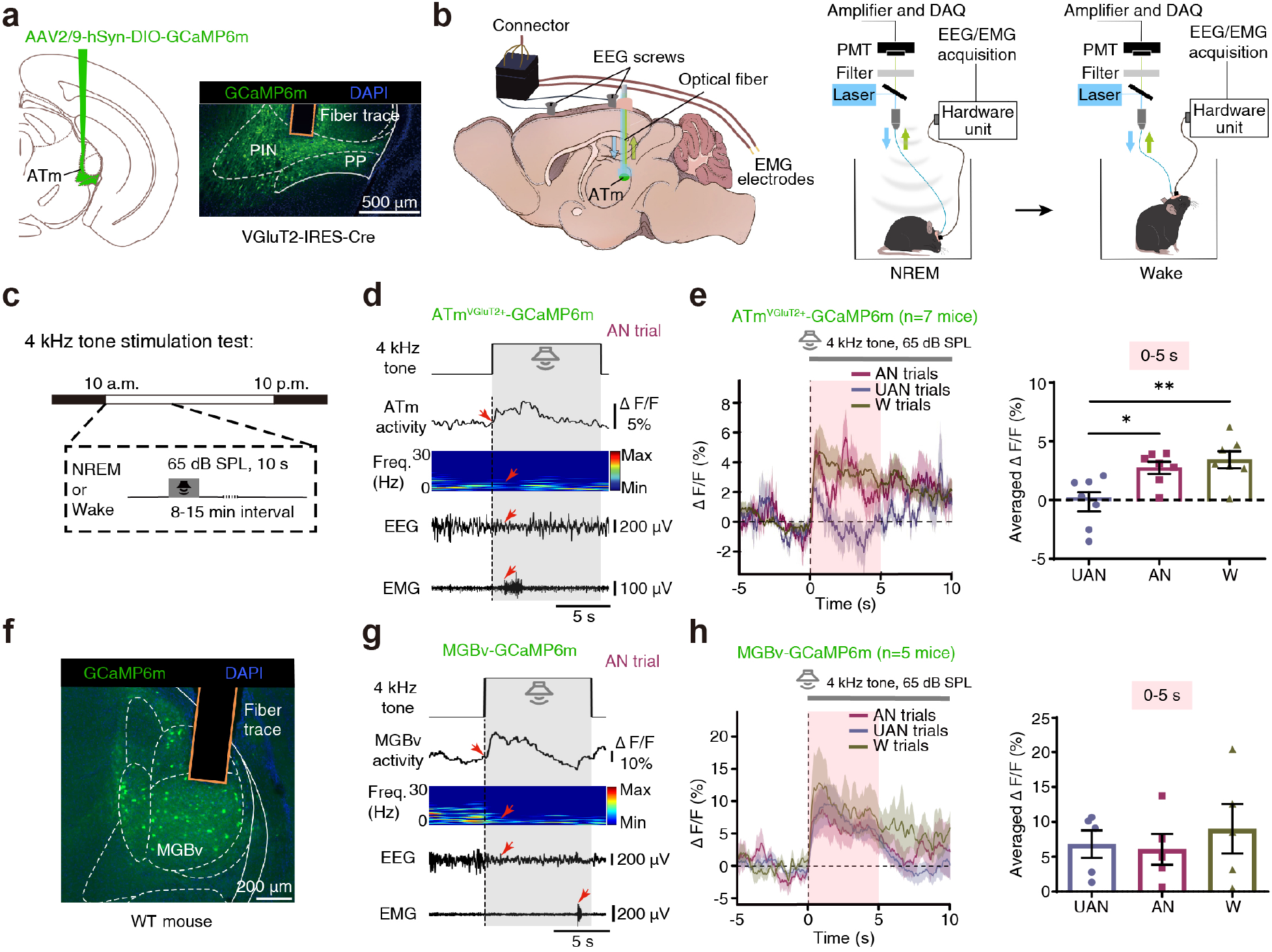
Auditory responsiveness of ATm^VGluT2+^ but not MGBv neuronal population correlates with sleep-wake states. **a.** Left: Schematic of virus injection in the ATm of VGluT2-IRES-Cre mouse for fiber photometry. Right: Expression of GCaMP6m in and fiber trace for recording from two of the ATm subnuclei populated by VGluT2+ neurons. Scale bar: 500 μm. **b.** Schematic design showing simultaneous monitoring of ATm^VGluT2+^ population-level Ca^2+^ activity (abbreviated as ATm^VGluT2+^ activity later on for convenience) induced by pure tone and recording of EEG/EMG signals in a freely-moving mouse across sleep-wake states. **c.** Schematic diagram illustrating the parametric protocol for presenting sound stimulation. A pure tone (4 kHz, 65 dB SPL, 10 s duration) was manually applied to mice with inter-stimulus-interval (ISI) of 8-15 minutes during light cycle (10:00 a.m.-10:00 p.m.). **d.** Tone-induced ATm^VGluT2+^ activity, EEG power spectrum, and EEG/EMG signals in an example awakened (AN; NREM-to-Wake) trial. Vertical dotted line indicates the onset of 4 kHz tone stimulation; Translucent gray area indicates the duration of tone; Red arrows from top to bottom indicate the onset of tone-induced ATm^VGluT2+^ activity, delta (0.5-4 Hz) power reduction, cortical activation, and behavioral activation, respectively. Scale bar: 5 seconds. **e.** Left: Averaged ATm^VGluT2+^ activity in AN, un-awakened (UAN) and wakefulness (W) trials. Vertical dotted line indicates the onset of stimulation; Translucent pink area indicates the time period (5 seconds) within which most tone-induced awakenings occurred. Right: Averaged ΔF/F values in the first 5 seconds after tone onset in three trial types. *n* = 7 mice, UAN vs. AN: *P* = 0.0209 (*t_6_* = 3.108); UAN vs. W: *P* = 0.0065 (*t_12_* = 3.285). **f.** Expression of AAV1-hSyn-GCaMP6m in and fiber trace for recording from the MGBv of a WT mouse. Scale bar: 200 μm. **g.** Tone-induced MGBv activity, EEG power spectrum, and EEG/EMG signals in an example AN trial. Scale bar: 5 seconds. **h.** Left: Averaged MGBv activity in AN, UAN and W trials. Right: Averaged ΔF/F values in the first 5 seconds after tone onset in three trial types. *n* = 5 mice. ATm, the medial sector of the auditory thalamus; PIN, the posterior intralaminar thalamic nuclei; PP, the peripeduncular nucleus; MGBv, the ventral division of the medial geniculate body. Paired or unpaired *t* test (two-tailed) was used in (e, h) for statistical analysis when data was normally distributed. Data are represented as mean (lines or bars) ± SEM (shaded regions or error bars); \**P* < 0.05, \*\**P* < 0.01. See also Supplementary Fig. 1 and Supplementary Table 1.

Three to four weeks after the surgery, we examined sensory evoked responses of ATm^VGluT2+^ neurons by presenting either auditory (4 kHz tone; 65 dB SPL, 10 s duration) or visual (blue light ^31^; 800 lx, 30 s duration) stimuli. Due to the relative rareness of REM sleep, sensory stimuli were delivered during either NREM sleep (N trials) or wakefulness (W trials) within the light cycle (Fig. 1c; Supplementary Fig. 1A). According to whether sleep-to-wake transition, which is signaled by the appearance of cortical activation (CA) ^32^ (See Methods for detailed procedure), occurred shortly after the presentation of sensory stimuli, NREM trials were further categorized into awakened NREM (AN) and un-awakened NREM (UAN) trials. Since tone- and blue light-induced cortical activation predominantly appeared in the first 5 s and 10 s after stimuli presentation, respectively, our data analysis focused on corresponding time windows. We found that although onset Ca^2+^ signals in response to sensory stimuli was reliably observed (Fig. 1d; Supplementary Fig. 1B), sensory evoked Ca^2+^ signals in UAN trials exhibited strong adaptation to continuous tone and blue light (Fig. 1e; Supplementary Fig. 1C, left panel), leading to significantly smaller signal increase when averaged across respective time window (Fig. 1e; Supplementary Fig. 1C, right panel). None of these phenomena was observed in VGluT2-Cre mice injected with AAV expressing eGFP in the ATm (Supplementary Fig. 1D). These data demonstrated that, exactly as we expected, the sensory responsiveness of ATm^VGluT2+^ neurons during NREM sleep closely correlated with the probability of sensory evoked sleep-to-wake transition.

Compared with ATm^VGluT2+^ neurons, how would those in the primary sensory thalamic nuclei, which relay stimulus feature-related information to the primary sensory cortices, respond to sensory stimuli during NREM sleep? Although a previous study conducted in the auditory cortex of Marmoset monkey suggested that the sound-driven responses of MGBv neurons at population level should not be significantly different across sleep-wake states ^16^, direct evidence remained lacking. Different from the prevalence of VGluT2+ neurons in the ATm, either VGluT1+ or VGluT2+ cells only represent a subset of excitatory neurons in the MGBv ^33^, which does not contain other subtype neurons ^34^. To record Ca^2+^ population signal from as many MGBv neurons as possible, we expressed GCaMP6m in MGBv neurons by injecting AAV1-hSyn-GCaMP6m into the MGBv of wild type (WT) mice followed by implanting optical fiber into the MGBv (Fig. 1f). As expected, MGBv Ca^2+^ population signal demonstrated strong onset responses to tonal stimuli in all trials (Fig. 1g). However, in contrast to what we observed in ATm^VGluT2+^ neurons, sound-evoked MGBv Ca^2+^ signal persisted through the designated time window (first 5 s after tone onset) in all three trial types (Fig. 1h, left panel), resulting in state-independent Ca^2+^ signal increase (Fig. 1h, right panel). To physiologically verify that the sound-driven Ca^2+^ signals were predominantly recorded from the MGBv but not the ATm, which surrounds the MGBv and is responsive to blue light, we also recorded from the MGBv while presenting blue light during NREM sleep. Clearly, Ca^2+^ signal did not show onset response to blue light (Supplementary Fig. 1E, left panel) and did not significantly increase within the designated time window (first 10 s after blue light onset) (Supplementary Fig. 1E, right panel). In control mice injected with eGFP-expressing AAV, neither tone nor blue light induced significant Ca^2+^ signal change in the MGBv (Supplementary Fig. 1F, G). Thus, distinct from ATm^VGluT2+^ neurons, MGBv neurons exhibited state-independent reliable population responses to auditory stimuli, echoing with the phenomena observed in the primary auditory cortex of marmoset monkey ^16^. Nevertheless, it is worth noting again that responses observed at population level do not necessarily reflect those at single neuron level.

### Photoactivating ATm^VGluT2+^ rather than lemniscal thalamic neurons induces rapid NREM sleep-to-wake transition

Based on the close correlation between sensory evoked responses of ATm^VGluT2+^ neurons and probability of awakening from NREM sleep, we inferred that photoactivating ATm^VGluT2+^ neurons could induce NREM sleep-to-wake transition. To test this hypothesis, we conducted the same set of surgeries in mice as we did for recording Ca^2+^ signals from the ATm, except that AAV2/9-EF1α-DIO-hChR2-eYFP was unilaterally injected instead for photoactivation (Fig. 2a, b, upper panel). After the mice recovered from surgery, laser stimuli with a duration of 20 s (473 nm, 10 Hz, 10 ms, 20 s pulse train) was delivered to ATm^VGluT2+^ neurons during light cycle (10 a.m. - 10 p.m.) at random time points between 10-15 minutes after the offset of last laser stimulation (Fig. 2b, bottom panel).

**Fig. 2.**
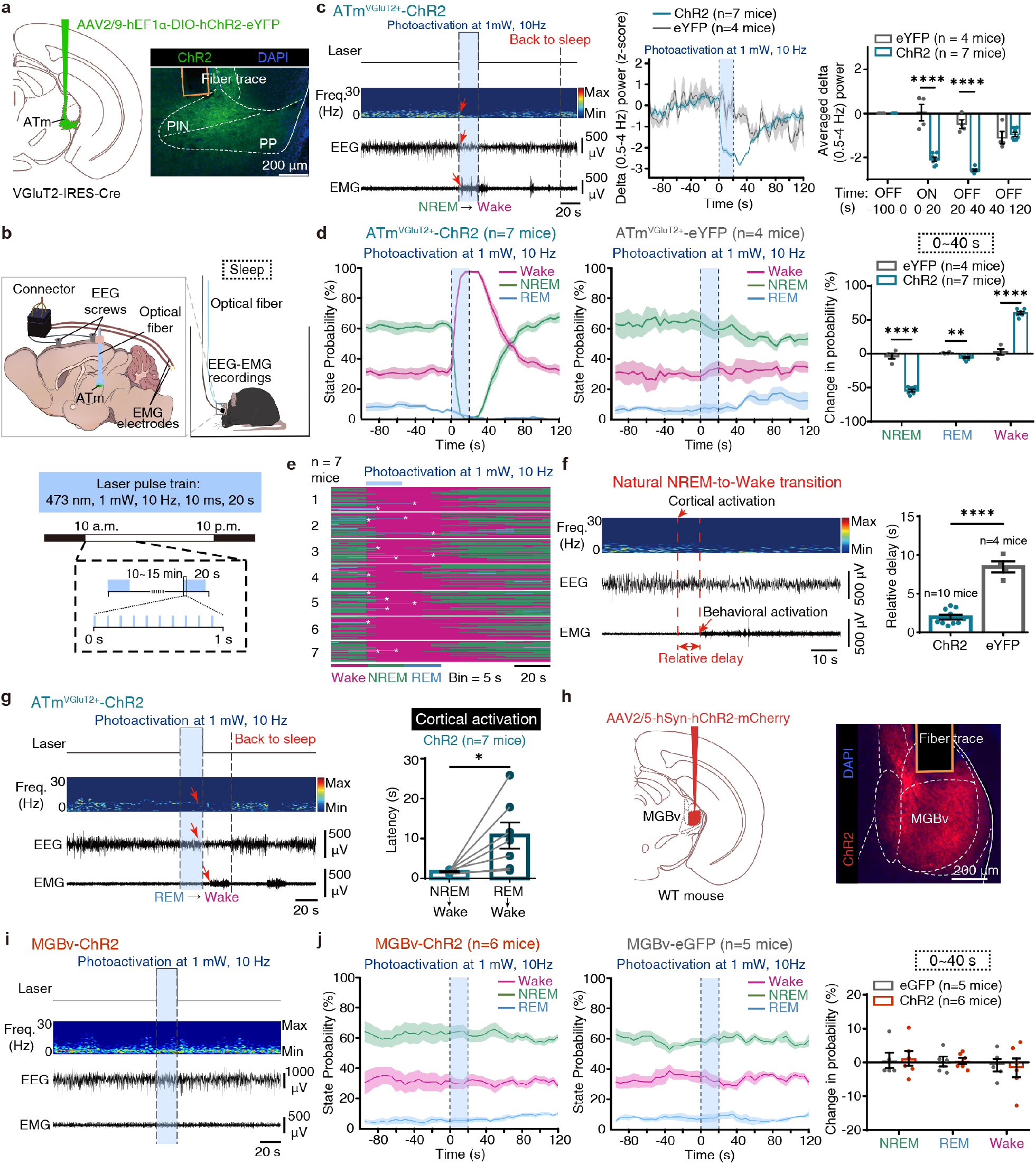
Photoactivation of ATm^VGluT2+^ rather than MGBv neurons induces rapid NREM-to-wake transition. **a.** Left: Schematic of virus injection in the ATm of a VGluT2-IRES-Cre mouse for photoactivation. Right: Representative expression of ChR2 and fiber trace for photoactivation in two of the ATm subnuclei populated by VGluT2+ neurons. Scale bar: 200 μm. **b.** Upper panel: Schematic design showing simultaneous photoactivation of ATm^VGluT2+^ neurons and recording of EEG/EMG signals in a freely-moving mouse. Lower panel: Schematic diagram illustrating the parametric protocol for photoactivation. Laser pulses (1 mW, 10 Hz, 10 ms, 20 s duration) were delivered during light cycle (10:00 a.m.-10:00 p.m.) with ISI of 10-15 minutes. **c.** Left: EEG power spectrum and EEG/EMG signals in a representative trial showing ATm^VGluT2+^ photoactivation-induced NREM-to-Wake transition. Translucent light-blue area indicates the duration of photoactivation; Red arrows from top to bottom indicate the onset of delta power reduction, cortical activation, and behavioral activation, respectively; The right most vertical dashed line indicates the time point at which the mouse reentered sleep. Scale bar: 20 seconds. Middle: Averaged delta power before, during and after ATm^VGluT2+^ photoactivation during NREM sleep in ChR2- (*n* = 7 mice) and eYFP- (*n* = 4 mice) mice. Translucent light-blue area indicates the duration of photoactivation. Right: Delta power values averaged in different time windows relative to photoactivation onset. ChR2 vs. eYFP: laser ON (0-20 s), *P* < 0.0001 (*t_9_* = 7.074); laser OFF (20-40 s), *P* < 0.0001(*t_9_* = 13.45). **d.** State probability before, during and after ATm^VGluT2+^ photoactivation in ChR2-(left) and eYFP-(middle) mice during light cycle. Right: Averaged changes in state probability within 40 seconds after photoactivation onset. NREM sleep, *P* < 0.0001 (*t_9_* = 12.4); REM sleep, *P* = 0.0022 (*t_9_* = 4.239); Wake, *P* < 0.0001 (*t_9_* = 13.93). **e.** Photoactivation trials in all 7 ATm^VGluT2+^-ChR2 mice. White asterisks indicate REM-to-Wake transitions. **f.** Left: A representative natural NREM-to-Wake transition indicated by changes in EEG power spectrum and EEG/EMG signals. Red arrows from top to bottom indicate the onset of cortical and behavioral activation, respectively. The time window between two vertical dashed lines is the relative delay between cortical and behavioral activation. Right: The relative delay in ChR2-mice (*n* = 10) following photoactivation onset and in eYFP-mice (*n* = 4). ChR2 vs. eYFP: *P* < 0.0001 (*t_12_* = 9.908). **g.** Left: EEG power spectrum and EEG/EMG signals in a representative trial showing photoactivation-induced REM-to-Wake transition. Right: The latency of photoactivation-induced NREM- or REM-to-Wake transition. NREM→Wake vs. REM→Wake: *n* = 7 mice, *P* = 0.0311 (*t_6_* = 2.803). **h.** Left: Schematic of virus injection in the MGBv of WT mouse. Right: Representative expression of ChR2 in the MGBv and fiber trace for photoactivation. Scale bar, 200 μm. **i.** EEG power spectrum and EEG/EMG signals in a representative MGBv-photoactivation trial. Scale bar: 20 seconds. **j.** State probability before, during and after MGBv photoactivation in ChR2- (left, *n* = 6 mice) and eGFP- (middle, *n* = 5 mice) mice during light cycle. Right: Averaged changes in state probability within 40 seconds after photoactivation onset. Paired *t* test (two-tailed) was used in (g), and unpaired *t* test (two-tailed) was used in (c, d, f, j) when data was normally distributed, otherwise Mann-Whitney *U* test was applied for statistical analysis. Data are represented as mean (lines or bars) ± SEM (shaded regions or error bars); \**P* < 0.05, \*\**P* < 0.01, \*\*\*\**P* < 0.0001. See also Supplementary Fig. 2 and Supplementary Table 1.

Strikingly, we found that the arousing effect of ATm photoactivation was extraordinarily strong even when laser power was as low as 1 mW. Specifically, photoactivating ATm^VGluT2+^ neurons during NREM sleep induced immediate (latency: ∼1.7 s) cortical activation (desynchronized EEG signal) and behavioral activation (desynchronized EEG signal coupled with elevated EMG signal; See Methods for detailed description) (Fig. 2c, left panel; 2g, right panel; Supplementary Movie 1). A significant decrease in EEG delta power (0.5-4 Hz), which is a measure of the intensity of NREM sleep, lasted for at least 40 s after the stimulation onset (Fig. 2c, middle and right panels), accompanied by a robust decrease and increase of NREM sleep and wake probability, respectively (Fig. 2d, left and right panels). This rapid and marked photoactivation-induced NREM sleep-to-wake transition was observed in vast majority of stimulation trials (Fig. 2e, Green lines). Continuous photoactivation of ATm^VGluT2+^ neurons yielded sleep-to-wake transition and shelter-seeking behavior (data not shown), indicating that the arousal induced by ATm activation was distinct from the microarousals spontaneously occur during sleep ^35^. Furthermore, compared with natural NREM sleep-to-wake transitions that are accompanied by behavioral activation, photoactivating ATm^VGluT2+^ neurons significantly shortened the relative delay between cortical and behavioral activation from ∼8 to 2 s (Fig. 2c, left panel; 2f), suggesting a behavior-activating role of ATm^VGluT2+^ neuronal activity in addition to its arousing effect. Interestingly, if photoactivation occurred during REM sleep, the time required for the mice to be awakened would be much longer (∼11 s) (Fig. 2g), although the decrease of REM sleep probability was still significant (Fig. 2d, left and right panels). In control mice injected with eYFP-expressing AAV, all the phenomena mentioned above were not observed (Fig. 2c, d, f; Supplementary Fig. 2A). Thus, photoactivating ATm^VGluT2+^ neurons can induce a rapid transition from sleep, NREM sleep in particular, to wakefulness both electrocortically and behaviorally.

Although our fiber photometry recordings across sleep-wake states from MGBv neurons strongly suggested that the lemniscal thalamus does not contribute to sensory evoked arousal (Fig. 1h), further evidences were still needed. To this end, we unilaterally injected AAV2/5-hSyn-hChR2-mCherry into the MGBv of WT mice for photoactivating MGBv neurons under different brain states (Fig. 2h). As expected, photoactivating MGBv neurons at a laser power of 1 mW did not induce NREM sleep-to-wake transition and did not change the probability of all three brain states (Fig. 2i, j, left and right panels; Supplementary Movie 2). In order to exclude the possibility that laser power was too low to induce sleep-to-wake transition, we increased laser power to 10 mW and still did not observe any arousing effect (Supplementary Fig. 2B, C). To confirm that neurons in the primary auditory cortex (Au1) were indeed activated by cortical projecting neurons in the MGBv during photoactivation, we simultaneously recorded from the ipsilateral Au1 by injecting RCaMP-expressing rAAV-hSyn-NES-jRGECO1a into the Au1 of one of the ChR2 mice (Supplementary Fig. 2D). Indeed, Au1 neurons demonstrated robust onset and facilitating responses to MGBv photoactivation, and the magnitude was positively correlated with laser power (Supplementary Fig. 2E). We further extended the same set of experiments to the dLGN, the primary visual thalamus (Supplementary Fig. 2F), and found that photoactivating dLGN at laser power of 1 or 10 mW did not induce any change in brain states as well (Supplementary Fig. 2G, H; Supplementary Movie 2). Based on these findings, we concluded that the primary sensory thalamic nuclei, the MGBv and dLGN in present study, and their cortical projections do not play a role in arousal.

### ATm^VGluT2+^ neuronal activity is indispensable for both blue light- and sound-induced awakening

We next determined whether and to what extent ATm^VGluT2+^ neuronal activity would be important for sensory-evoked awakening. To address this question, we unilaterally injected AAV2/9-CAG-DIO-GtACR1-GFP into the ATm of VGluT2-IRES-Cre mice to photoinhibit ATm^VGluT2+^ neurons^36, 37^ (Fig. 3a). To match with the unilateral photoactivation experiment described above, we injected AAV2/9-CAG-DIO-taCasp3, which was mixed with AAV2/9-CAG-DIO-GtACR1-GFP for control purpose, into the ATm of contralateral side to ablate ATm^VGluT2+^ neurons ^38^. For each mouse, once baseline tests for sensory evoked awakening was completed, the same set of tests combined with photoinhibition were conducted at least 2 days later (Fig. 3b; see Methods for detailed procedure). Considering that photoinhibition is less efficient than photoactivation in terms of manipulating neural activity ^39^, we opted to turn on the laser 20 s before the delivery of sound or blue light during NREM sleep (Fig. 3b, bottom panel).

**Fig. 3.**
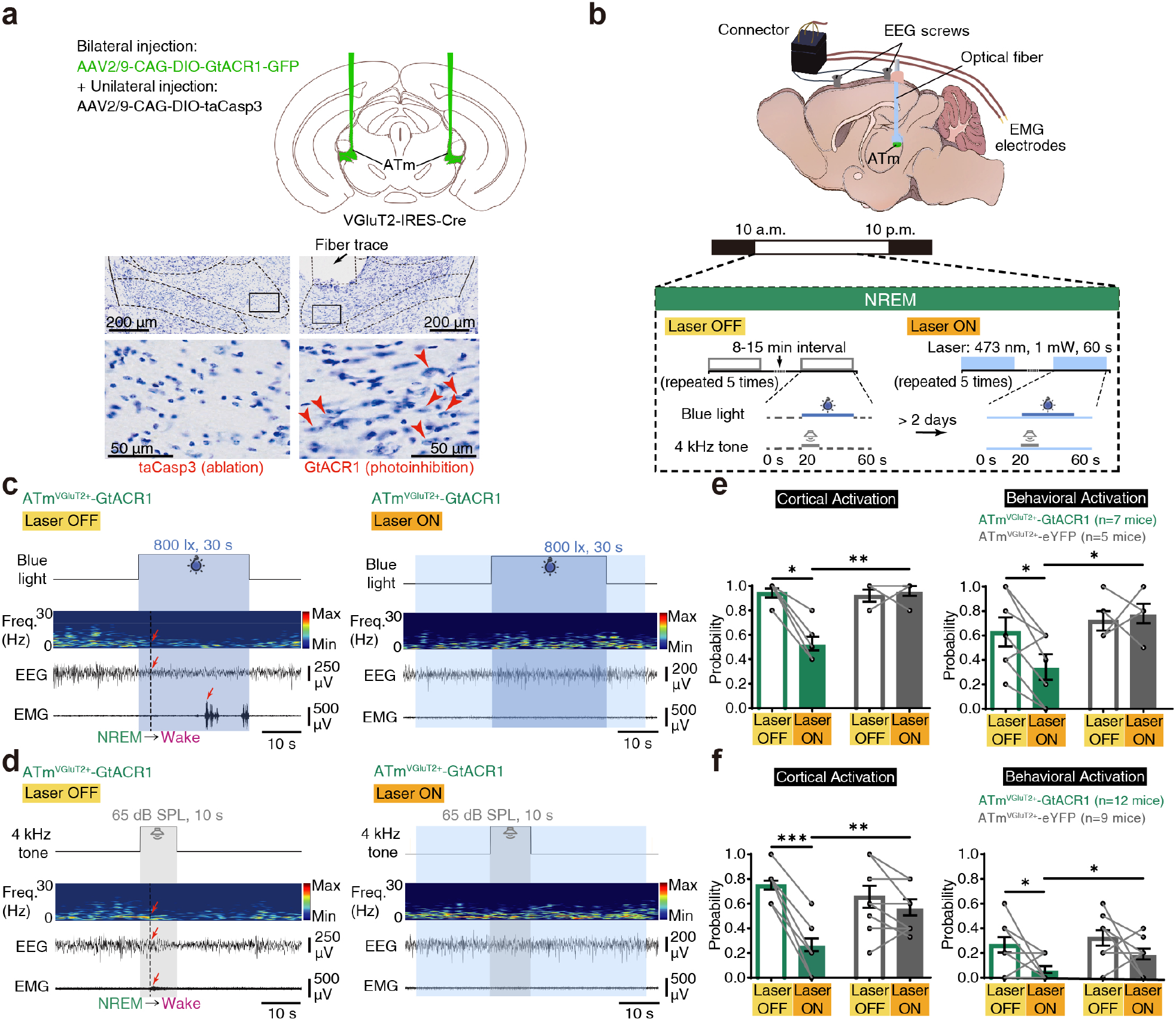
ATm^VGluT2+^ neuronal activity is indispensable for both blue light- and sound-induced awakening. **a.** Upper panel: Schematic of virus injection in the ATm of VGluT2-IRES-Cre mouse for photoinhibition. Middle- and bottom-left panels: Representative Nissl staining showing unilateral ablation of ATm^VGluT2+^ neurons by the expression of taCasp3. Middle- and bottom-right panels: Intact ATm^VGluT2+^ neurons (indicated by red arrow heads) on the other side. Scale bars: 200 μm (middle) or 50 μm (bottom). **b.** Upper panel: Schematic design showing simultaneous photoinhibition of ATm^VGluT2+^ neurons and recording of EEG/EMG signals in a freely-moving mouse. Lower panel: Schematic diagram illustrating the parametric protocol for the delivery of laser stimulus (473 nm, 1 mW, 60 s) and blue light (800 lx, 30 s) or 4 kHz tone (65 dB SPL, 10 s). The baseline tests (Laser OFF trials) were conducted more than 2 days before photoinhibition tests (Laser ON trials). **c-d.** Blue light- or tone-induced changes in EEG power spectrum and EEG/EMG signals in representative Laser OFF (left) and Laser ON (right) trials. **e.** Probability of blue light-induced cortical (left) and behavioral (right) activation in GtACR1- (Green, *n* = 7 mice) and eYFP- (Gray, *n* = 5 mice) mice in Laser OFF and ON trials. Left: GtACR1, Laser OFF vs. Laser ON, *P* = 0.0156 (*W* = −28); Laser ON, GtACR1 vs. eYFP, *P* = 0.0051 (*U* = 0.5). Right: GtACR1, Laser OFF vs. Laser ON, *P* = 0.0249 (*t_6_* = 2.970); Laser ON, GtACR1 vs. eYFP, *P* = 0.0115 (*t_10_* = 3.086). **f.** Probability of tone-induced cortical (left) and behavioral (right) activation in GtACR1- (Green, *n* = 12 mice) and eYFP- (Gray, *n* = 9 mice) mice in Laser OFF and ON trials. Left: GtACR1, Laser OFF vs. Laser ON, *P* = 0.0005 (*W* = −78); Laser ON, GtACR1 vs. eYFP, *P* = 0.0015 (*t_19_* = 3.715); Right: GtACR1, Laser OFF vs. Laser ON, *P* = 0.0391 (*W* = −31); Laser ON, GtACR1 vs. eYFP, *P* = 0.0412 (*U* = 26). Paired or unpaired *t* test (two-tailed) was used in (e, f) when data was normally distributed, otherwise Wilcoxon matched-pairs signed rank test or Mann-Whitney *U* test was applied for statistical analysis. Data are represented as mean (bars) ± SEM (error bars); \**P* < 0.05, \*\**P* < 0.01, \*\*\**P* < 0.001. See also Supplementary Fig. 3 and Supplementary Table 1.

We found that, compared with GtACR1 mice that were not photoinhibited (pre-inhibition), mice with inhibited ATm^VGluT2+^ neurons were difficult to be awakened from NREM sleep by both blue light and tone (Fig. 3c, d; Supplementary Movie 3). Photoinhibition decreased the probability of blue light- and tone-induced cortical activation from 94% and 75% to 53% and 27%, respectively, and the reduction was larger than 40% of pre-inhibition awakening probability (Fig. 3e, f, left panel). Thus, sensory evoked awakening occurred at chance level or even lower probability once ATm^VGluT2+^ neurons were photoinhibited. In terms of blue light- and tone-induced behavioral activation, photoinhibiting ATm^VGluT2+^ neurons lowered its probability from 63% and 27% to 34% and 7%, respectively, and the reduction was larger than 45% of pre-inhibition probability (Fig. 3e, f, right panel). These findings strongly indicated that, under our experimental conditions, sensory evoked activity of ATm^VGluT2+^ neurons was indispensable for blue light- and tone-induced arousal. All the effects observed in photoinhibited GtACR1 mice were not observed in control mice expressing eYFP in ATm^VGluT2+^ neurons (Fig. 3e, f).

We further confirmed that ATm indeed plays an important role in mediating sensory evoked arousal by photoactivating visual and auditory input pathways separately. We unilaterally injected AAV2/9-hEF1α-DIO-hChR2-eYFP into either the cortex of the inferior colliculus (ICx) or the optic and deeper layers of the superior colliculus (SO/dSC), which provide ascending auditory or visual inputs to the ATm ^21^, of VGluT2-IRES-Cre mice, followed by optical fiber implantation targeting the ATm (Supplementary Fig. 3A). Neurons expressing VGluT2 is the major type of excitatory neuron in both dorsal midbrain regions ^33, 40^. Similar to the effect of stimulating ATm^VGluT2+^ neurons, photoactivating either SO/dSC^VGluT2+^-ATm or ICx^VGluT2+^-ATm pathway induced immediate cortical and behavioral activation (Supplementary Fig. 3B). The probability of NREM sleep and wake markedly decreased and increased following photoactivation onset, respectively, but that of REM sleep was unchanged (Supplementary Fig. 3C, D, left and right panels). In control mice injected with eGFP-expressing AAV, the laser stimulation did not cause any significant change in state probability (Supplementary Fig. 3C, D, middle panel).

### Photoactivating ATm^VGluT2+^-TeA/ECT^Pyr^ pathway induces NREM sleep-to-wake transition

To gain insight into the downstream neural circuits involved in mediating sensory evoked arousal, we next examined the efferent targets of ATm^VGluT2+^ neurons. Like other sensory thalamic nuclei, the ATm also makes cortical projections, which mainly concentrate in the TeA and more ventrally located ECT ^21^, which are both sensory association cortices. Two recent studies revealed that normal TeA activity is critical for social behaviors, such as social interactions between adult mice ^41^ and auditory-driven pup-retrieval behavior ^42^ that putatively require a certain level of arousal. Of note, neuropathological and functional abnormalities are particularly prominent in the TeA, mainly affecting pyramidal (Pyr) neurons, of autistic children that exhibit both emotional overreaction and hyperarousal to sensory stimulation ^43, 44^. These studies suggested to us that TeA/ECT^Pyr^ neuronal activity might simultaneously promote arousal while playing its role in social behaviors, and inspired us to hypothesize that the axonal fibers from the ATm might innervate TeA/ECT^Pyr^ neurons, thereby contributing to the sensory evoked arousal mediated by the ATm.

To test the hypothesis mentioned above, we first injected scAAV2/1-hSyn-Cre into the ATm of WT mice to transsynaptically express Cre in downstream TeA/ECT neurons, and injected AAV2/9-hEF1α-DIO-mCherry into the TeA/ECT region to enable Cre-dependent expression of mCherry (Supplementary Fig. 4A). Exactly as expected, this transsynaptic tracing approach predominantly labeled TeA/ECT^Pyr^ neurons (Supplementary Fig. 4B, top-right two panels), along with a small fraction of GABAergic neurons (Supplementary Fig. 4B, bottom-left two panels), which putatively provide feedforward inhibition. TeA/ECT^Pyr^ neurons receiving ATm inputs were mainly located in layer V followed by layer II-III (Supplementary Fig. 4B, top-left panel). Since pyramidal neurons are the major source of long-range cortico-cortical/subcortical projections, we further injected AAV1-CAG-GFP into the TeA/ECT of WT mice (Supplementary Fig. 4C). We observed dense axonal fibers in but not limited to the nucleus accumbens ^45, 46, 47^ and claustrum ^48, 49^ (Supplementary Fig. 4D), which are critically involved in the regulation of arousal and consciousness. This projection pattern further suggested that the ATm^VGluT2+^-TeA/ECT^Pyr^ pathway might have contributed to the sensory evoked arousal observed in our preceding experiments.

Toward this end, we first tested whether selectively photoactivating the ATm^VGluT2+^- TeA/ECT^Pyr^ pathway could induce arousal. We unilaterally injected AAV2/9-hEf1α-DIO-hChR2-eYFP into the ATm of VGluT2-IRES-Cre mice (Fig. 4a), then we fixed an optical fiber to the thinned skull above the TeA/ECT perpendicularly, which enables transcranial photoactivation of ChR2-expressing axons from the ATm (473 nm, 10 ms, 10 Hz, 15-30 mW, 20 s). Laser stimulation of ATm^VGluT2+^ axons in the TeA/ECT during NREM sleep induced cortical activation in around 64% of trials (in less than 10 s) followed by behavioral activation (Fig. 4b; Supplementary Fig. 4E; Supplementary Movie 4), as shown by a significant decrease in EEG delta power that lasted for at least 40 s after the stimulation onset (Fig. 4c) and by a significant decrease and increase in NREM sleep and wake probability, respectively (Fig. 4d, left and right panels). The probability of REM sleep was not significantly changed. The arousing effects of photoactivation were not observed in control mice (Fig. 4d, middle and right panels; Supplementary Fig. 4F). Thus, selective photoactivation of the ATm^VGluT2+^-TeA/ECT^Pyr^ pathway was indeed able to induce NREM sleep-to-wake transition.

**Fig. 4.**
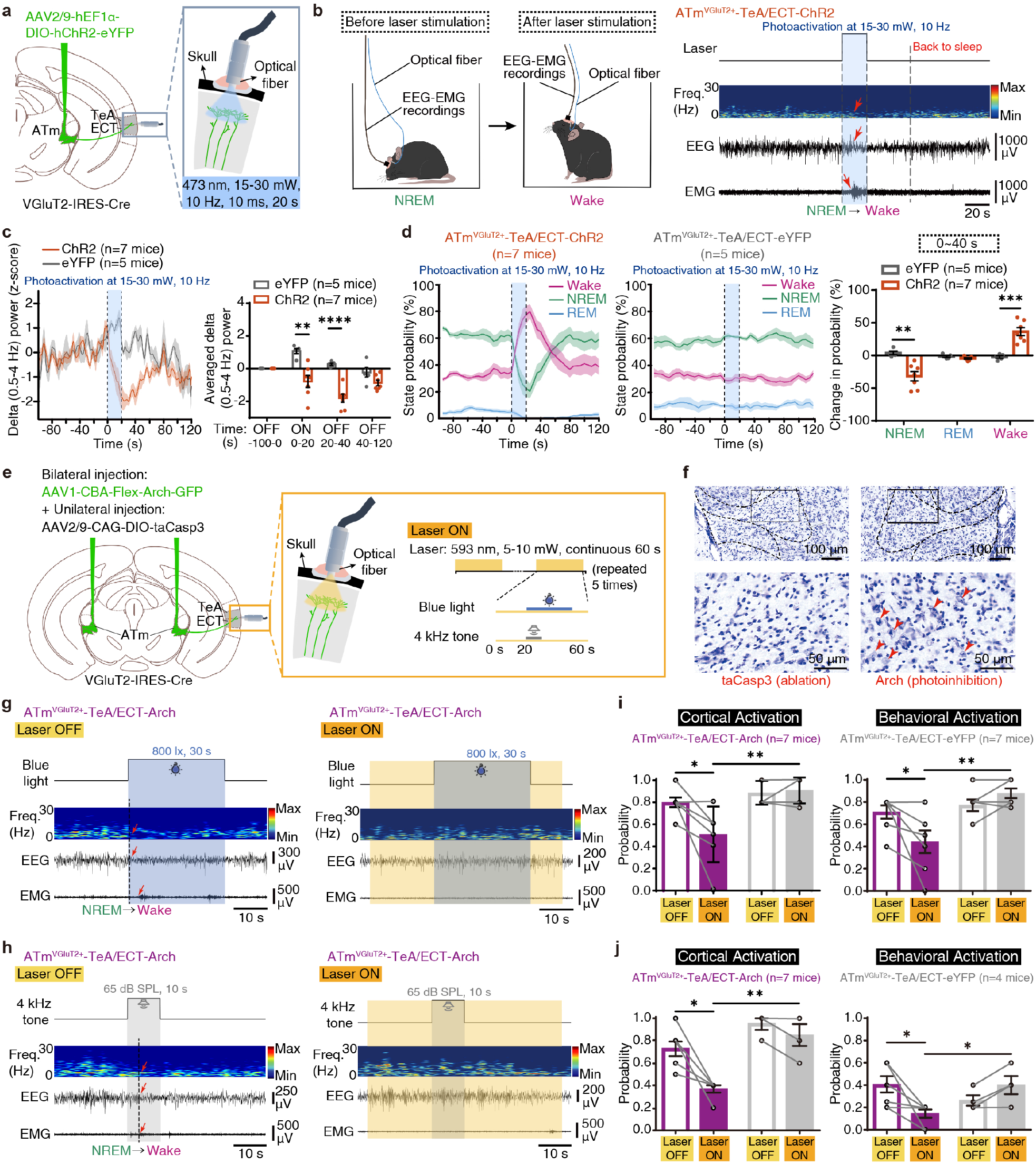
The ATm^VGluT2+^-TeA/ECT pathway contributes to both blue light- and sound-induced awakening. **a.** Schematic of experimental set-up for transcranial photoactivation of ATm^VGluT2+^ axonal fibers in the TeA/ECT. Inset: An optical fiber perpendicularly fixed to the thinned skull over the TeA/ECT region. **b.** Left: Diagram of NREM-to-Wake transition induced by ATm^VGluT2+^-TeA/ECT pathway activation. Right: EEG power spectrum and EEG/EMG signals in a representative trial showing photoactivation-induced NREM-to-Wake transition. Translucent light-blue area indicates the duration of photoactivation; Red arrows from top to bottom indicate the onset of delta power reduction, cortical activation, and behavioral activation, respectively. Scale bar: 20 seconds. **c.** Left: Averaged delta power before, during and after ATm^VGluT2+^ axonal photoactivation during NREM sleep in ChR2- (*n* = 7 mice) and eYFP- (*n* = 5 mice) mice. Translucent light-blue area indicates the duration of photoactivation. Right: Delta power values averaged in different time windows relative to photoactivation onset. ChR2 vs. eYFP: laser ON (0-20 s), *P* = 0.0022 (*t_10_* = 4.085); laser OFF (20-40 s), *P* < 0.0001 (*t_10_* = 6.722). **d.** State probability before, during and after ATm^VGluT2+^ axonal photoactivation in ChR2- (left) and eYFP- (middle) mice during light cycle. Right: Averaged changes in state probability within 40 seconds after photoactivation onset. NREM sleep, *P* = 0.0023 (*t_10_* = 4.069); Wake, *P* = 0.0004 (*t_10_* = 5.209). **e.** Left: Schematic of experimental set-up for transcranial photoinhibition of ATm^VGluT2+^ axonal fibers in the TeA/ECT. Inset: Schematic diagram illustrating the parametric protocol for the delivery of laser stimulus (593 nm, 5-10 mW, 60 s) and blue light or 4 kHz tone. **f.** Left two panels: Representative Nissl staining showing unilateral ablation of ATm^VGluT2+^ neurons by the expression of taCasp3. Right: Intact ATm^VGluT2+^ neurons (indicated by red arrow heads) on the other side. Scale bar: 100 μm (upper panels) and 50 μm (lower panels). **g-h.** Left: Blue light- or tone-induced changes in EEG power spectrum and EEG/EMG signals in representative Laser OFF (left) and Laser ON (right) trials. **i.** Probability of blue light-induced cortical (left) and behavioral (right) activation in Arch- (Purple, *n* = 7 mice) and eYFP- (Gray, *n* = 7 mice) mice in Laser OFF and ON trials. Left: Arch, Laser OFF vs. Laser ON, *P* = 0.0313 (*W* = −21); Laser ON, Arch vs. eYFP, *P* = 0.0023 (*U* = 2). Right: Arch, Laser OFF vs. Laser ON, *P* = 0.0313 (*W* = −21); Laser ON, Arch vs. eYFP, *P* = 0.0029 (*U* = 2.5). **j.** Probability of tone-induced cortical (left) and behavioral (right) activation in Arch- (Purple, *n* = 7 mice) and eYFP- (Gray, *n* = 4 mice) mice in Laser OFF and ON trials. Left: Arch, Laser OFF vs. Laser ON, *P* = 0.0156 (*W* = −28); Laser ON, Arch vs. eYFP, *P* = 0.0030 (*U* = 0). Right: Arch, Laser OFF vs. Laser ON, *P* = 0.0313 (*W* = −21); Laser ON, Arch vs. eYFP, *P* = 0.0182 (*U* = 2.5). TeA, temporal association cortex; ECT, ectorhinal cortex. Unpaired *t* test (two-tailed) was used in (c, d), and paired *t* test (two-tailed) was used in (i, j) when data was normally distributed, otherwise Mann-Whitney *U* test was applied for statistical analysis. Data are represented as mean (lines or bars) ± SEM (shaded regions or error bars); \**P* < 0.05, \*\**P* < 0.01, \*\*\**P* < 0.001, \*\*\*\**P* < 0.0001. See also Supplementary Fig. 4-5 and Supplementary Table 1.

### The ATm^VGluT2+^-TeA/ECT^Pyr^ pathway contributes to both blue light- and sound-induced awakening

We next determined to what extent the ATm^VGluT2+^-TeA/ECT^Pyr^ pathway would be required for sensory evoked awakening by selectively photoinhibiting ATm^VGluT2+^ axons in the TeA/ECT. Briefly, AAV1-CBA-Flex-Arch-GFP was bilaterally injected into VGluT2-IRES-Cre mice targeting the ATm (Fig. 4e, left panel). Subsequently, AAV2/9-CAG-DIO-taCasp3 was unilaterally injected into the ATm to ablate VGluT2+ neurons (Fig. 4e, f). Then, on the other side, an optical fiber was fixed to the skull in the same way as described earlier to enable transcranial photoinhibition of Arch-expressing axons from the ATm (593 nm, 5-10 mW, continuous) (Fig. 4e, right panel). EEG and EMG recording electrodes were also implanted as described earlier. We found that photoinhibition of ATm^VGluT2+^-TeA/ECT^Pyr^ pathway effectively attenuated the arousing effect of both blue light and tone (Fig. 4g, h). The probability of blue light- and tone-induced cortical activation was reduced from 80% and 72% to 50% and 37% (Fig. 4i, j, left panel), and that of behavioral activation was decreased from 71% and 40% to 44% and 14% (Fig. 4i, j, right panel), respectively. Again, the eYFP control group did not show any significant effects of photoinhibition (Fig. 4i, j). Thus, the reduction of awakening probability resulting from transcranial axonal inhibition was larger than 35% of pre-inhibition probability in all circumstances, indicating an indispensable contribution of the ATm^VGluT2+^-TeA/ECT^Pyr^ pathway to both blue light- and tone-induced awakening.

Compared with the effects produced by direct manipulation of ATm^VGluT2+^ neuronal activity, those produced by the manipulation of the ATm^VGluT2+^-TeA/ECT^Pyr^ pathway were weaker, suggesting that other neural pathways arising from the ATm might also contribute to sensory evoked arousal. Indeed, the hypothalamus and striatum, which are involved in brain state modulation ^50, 51^, are also efferent targets of ATm^VGluT2+^ neurons ^21^. To test the potential role of hypothalamic and striatal projections arising from the ATm, the same set of surgical procedures as those done for investigating ATm^VGluT2+^-TeA/ECT^Pyr^ pathway were performed except that optical fiber was implanted targeting either the ventromedial hypothalamus (VMH) or the tail of the striatum (TS), which contains dense ATm^VGluT2+^ axonal fibers (Supplementary Fig. 5A). Only photoinhibition was applied to determine the contribution of these two pathways. Interestingly, although photoinhibiting the ATm^VGluT2+^-VMH pathway significantly reduced the probability of blue light-induced cortical and behavioral activation (Supplementary Fig. 5B), it had no effect on tone-induced awakening (Supplementary Fig. 5C). Conversely, photoinhibiting the ATm^VGluT2+^- TS pathway did not decrease the probability of blue light-induced arousal (Supplementary Fig. 5D), but it made the occurrence of tone-induced cortical activation more difficult (Supplementary Fig. 5E). These findings suggested that, at least under our experimental conditions, the ATm^VGluT2+^-VMH or -TS pathway contributes to sensory evoked arousal in a modality-specific way. Thereby, the ATm^VGluT2+^-TeA/ECT^Pyr^ pathway is the one that plays a modality-independent critical role in mediating sensory evoked arousal.

### ATm^VGluT2+^ neuronal activity promotes arousal associated with defensive emotion in awake behaving mice

Our preceding results demonstrated that ATm^VGluT2+^ neuronal activity is indispensable for alerting auditory and visual stimuli-induced sleep-to-wake transition, which is critical for animal’s survival. Nevertheless, would these neurons also play an important role in mediating alerting stimuli-induced brain and behavioral state transition in awake behaving mice? To address this question, AAV2/9-EF1α-DIO-hChR2-eYFP was bilaterally injected into the ATm followed by bilateral optical fiber implantation for optogenetic excitation (Fig. 5a). We first used pupillometry to measure the arousal level of awake head-fixed mice voluntarily navigating a cylindrical treadmill ^52^ (Fig. 5b, left panel). We found that the diameter of pupil started to increase soon after the onset of photoactivation and continued to increase over the duration of ATm^VGluT2+^ excitation (Fig. 5b, right panel). This phenomenon was observed in most mice and trials regardless of the locomotion state at photoactivation onset (Fig. 5c, left panel), yielding significantly enlarged pupil by the end of photoactivation (Fig. 5c, right panel). These data indicate that ATm^VGluT2+^ neuronal activity is able to promote arousal in awake behaving mice as well.

**Fig. 5.**
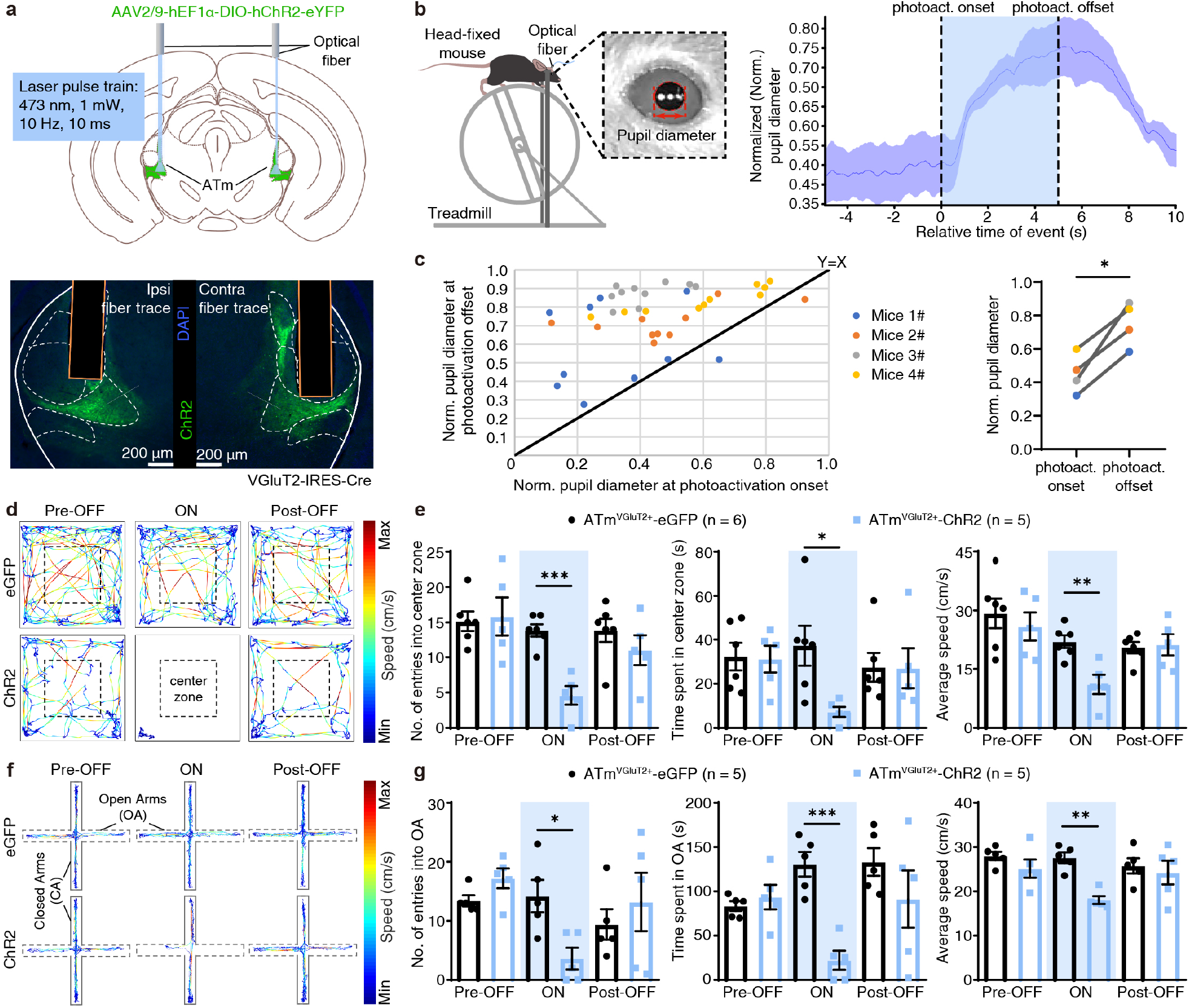
ATm^VGluT2+^ neuronal activity promotes arousal associated with defensive emotion in awake behaving mice. **a.** Upper panel: Schematic of virus injection and optical fiber implantation in the ATm of VGluT2-IRES-Cre mouse for photoactivation. Lower panel: Representative expression of ChR2 and fiber trace for photoactivation in the ATm. Scale bar: 200 μm. **b.** Left: Schematic design illustrating the experimental set-up for pupillometry in head-fixed awake behaving mouse. Right: Representative normalized (Norm.) pupil diameter of ChR2 mice before (−5-0 s), during (0-5 s) and after (5-10 s) photoactivation. **c.** Left: Norm. pupil diameter at photoactivation onset vs. offset. Each dot represents one photoactivation trial. Each color represents a different mouse. Right: Averaged Norm. pupil diameter at photoactivation onset and offset. Onset vs. offset (*n* = 4 mice): *P* = 0.0118 (*t_3_* = 5.511). **d.** Representative movement traces of eGFP- (upper row) and ChR2- (lower row) mice in a schematic open field (OF) before (left column, Pre-OFF), during (middle column, ON) and after (right column, Post-OFF) photoactivation. Dashed line square marks the center zone. The color gradient indicates the magnitude of movement speed in OF. **e.** The number of entries into the center zone (left), time spent in the center zone (middle), and average speed (right) of ChR2- (Sky blue, n = 5 mice) and eGFP- (Black, n = 6 mice) mice. eGFP (ON) vs. ChR2 (ON), left to right, *P* = 0.0002 (*t_9_* = 5.904); *P* = 0.0172 (*t_9_* = 2.914); *P* = 0.0041 (*t_9_* = 3.817). **f.** Representative movement traces of eGFP (upper row) and ChR2 (lower row) mice in a schematic elevated plus maze (EPM) before (left column, Pre-OFF), during (middle column, ON) and after (right column, Post-OFF) photoactivation. Solid lines represent closed arms (CA); dashed lines represent open arms (OA). The color gradient indicates the magnitude of movement speed in EPM. **g.** Number of entries into the OA (left), time spent in the OA (middle), and average speed (right) of ChR2- (Sky blue, n = 5 mice) and eGFP- (Black, n = 5 mice) mice. eGFP (ON) vs. ChR2 (ON), left to right, *P* = 0.0121 (*t_8_* = 3.225); *P* = 0.0003 (*t_8_* = 6.180); *P* = 0.0079 (*U* = 0). Paired *t* test (two-tailed) was used in (c), and unpaired *t* test (two-tailed) was used in (e, g) when data was normally distributed, otherwise Mann-Whitney *U* test was applied for statistical analysis. Data are represented as mean (lines or bars) ± SEM (shaded regions or error bars); \**P* < 0.05, \*\**P* < 0.01, \*\*\**P* < 0.001. See also Supplementary Fig. 6 and Supplementary Table 1.

Brain state at any given time can be thought of as a point in a high-dimensional space ^1^. Therefore, arousal is associated with other dimensions of brain state such as emotion depending on the activity of relevant neural circuits. Since ATm^VGluT2+^ neurons innervate multiple brain regions critically contributing to defensive emotional state ^21, 53^, we decided to determine whether the ATm^VGluT2+^-mediated arousal is associated with emotion of negative valence. We first conducted open field (OF) test and found that photoactivating ATm^VGluT2+^ neurons led to significant reduction in both the number of entries into and time spent in the center zone (Fig. 5d, bottom panel; 5e, left two panels). The average speed was also significantly decreased because the mice often stayed in one of the corners for a long time (Fig. 5e, right panel). These results suggested an anxiogenic effect of ATm^VGluT2+^ excitation. We further confirmed this conclusion by conducting elevated plus maze (EPM) test, in which the number of entries into and the time spent in the open-arm and average speed were all significantly reduced by ATm^VGluT2+^ excitation (Fig. 5f, g). We also exploited real-time place avoidance (RTPA) behavioral paradigm (Supplementary Fig. 6A), in which the number of entries into, the time spent in and average speed in the chamber paired with laser stimulation were significantly decreased (Supplementary Fig. 6B), suggesting the induction of fear or anxiety. All the stimulation effects mentioned above were not observed in control mice expressing eGFP rather than ChR2 in ATm^VGluT2+^ neurons (Fig. 5d-g, Supplementary Fig. 6). Taken together, these data indicate that the arousal promoted by ATm^VGluT2+^ neuronal activity was associated with enhanced negative emotion, consistent with the connectivity pattern of these neurons ^21^.

### ATm^VGluT2+^ neuronal activity is required for noise-triggered escape behavior

Since it is well known that brain state has profound impact on animal behavior ^1^, the activity of ATm^VGluT2+^ neurons might play an important role in triggering the expression of innate defensive behaviors when behaving mice are confronted with alerting stimuli. To test this hypothesis, we injected scAAV2/1-hSyn-Cre into the ICx, the non-lemniscal auditory midbrain, to transsynaptically express Cre in ATm neurons receiving ICx inputs, and injected AAV2/9-CMV-DIO-GFP-2A-TeNT into the ATm to block the synaptic transmission of Cre-expressing neurons ^54^ (Fig. 6a). Thereafter, we used an adapted behavioral assay (see Methods) to determine the importance of ATm^VGluT2+^ neurons receiving auditory inputs for sudden noise-provoked escape behavior. In this assay, a white noise (10 s) at 75 dB SPL was applied from the top once the mice entered the center zone of an open field (Fig. 6b). In response to this suddenly presented noise, control (eYFP) mice immediately fled towards a shelter nest placed in the corner and hid in there for about 18 seconds (eYFP: 18.3 ± 0.6 s), and their escape speed peaked at around 1.5 s (Fig. 6c, left panel), resulting in higher motion speed within 2 s after noise onset (Fig. 6d). In sharp contrast, TeNT mice did not behaviorally respond to the noise stimulation (Fig. 6c-e). In other words, they did not show increased motion speed during and after noise presentation, and did not spend as much time in the shelter (TeNT: 5.4 ± 4.3 s; Fig. 6d, f). Therefore, blocking the signaling of ATm^VGluT2+^ neurons receiving auditory inputs abolished sudden noise-induced escape behavior. Taken together, these data suggest that the arousal and negative emotion induced by ATm^VGluT2+^ neuronal activity are required for alerting stimuli to trigger the expression of innate defensive behaviors.

**Fig. 6.**
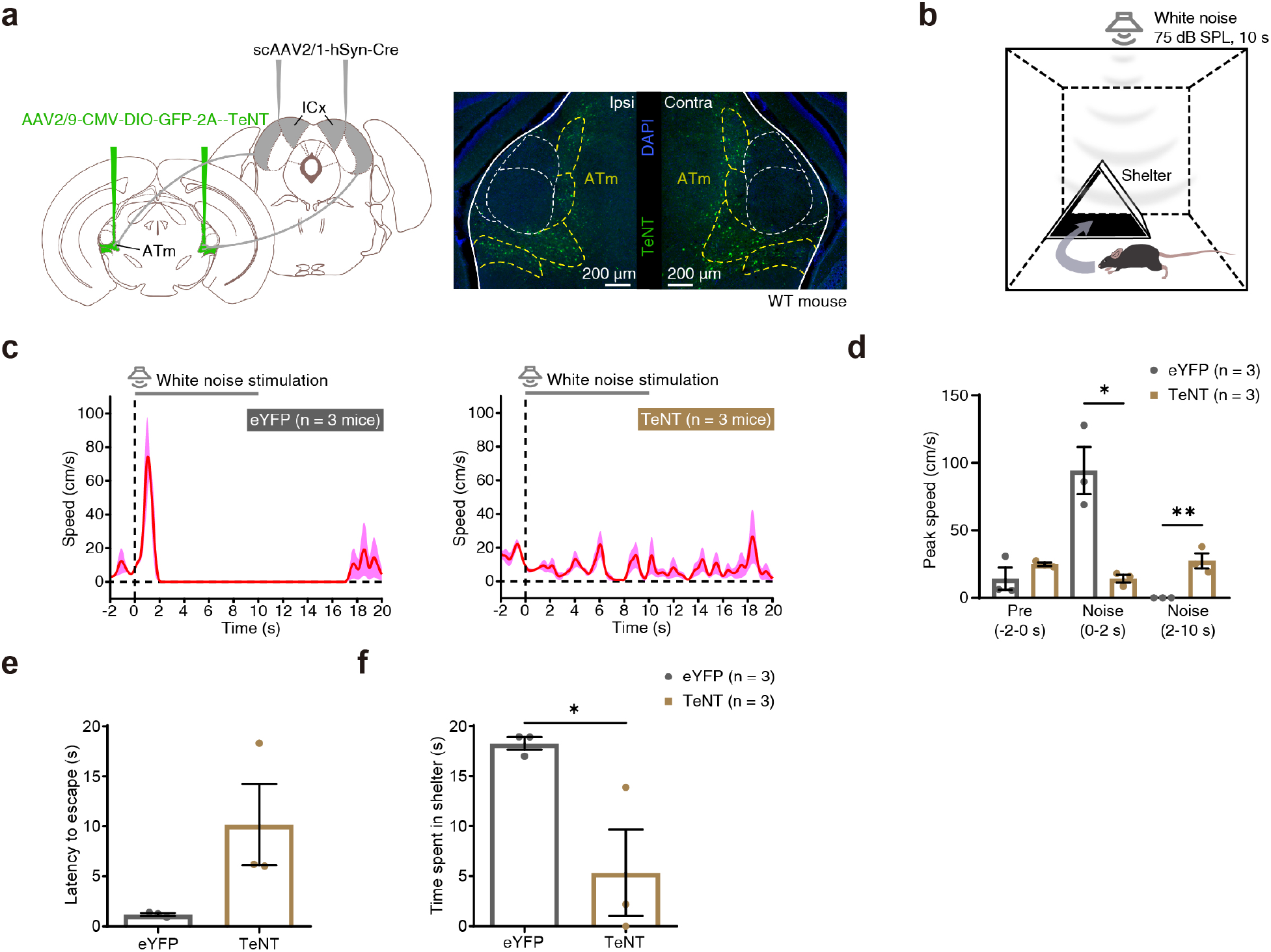
ATm^VGluT2+^ neuronal activity is required for noise-triggered escape behavior. **a.** Diagram showing virus injection strategy (upper panel) enabling anterograde trans-monosynaptic expression of both tetanus toxin (TeNT) and GFP in ATm neurons receiving ICx inputs (lower panel). Scale bar: 200 μm. **b.** Schematic of experimental set-up for noise-induced escape behavior. **c.** Averaged speed of eYFP- (upper panel, *n* = 3 mice) and TeNT- (lower panel, *n* = 3 mice) mice in the chamber shown in (b). Horizontal solid gray and lines indicate the duration of white noise stimulation. Vertical dotted black lines indicate the onset of white noise. **d.** Peak speed of mice before (−2-0 s) and during (0-2 s; 2-10 s) the application of white noise. eYFP (*n* = 3 mice) vs. TeNT (*n* = 3 mice): 0-2 s, *P* = 0.0106 (*t_4_* = 4.529); 2-10 s, *P* = 0.0078 (*t_4_* = 4.935). **e.** Latency to escape after the onset of noise representation in eYFP- (n = 3 mice) and TeNT- (n = 3 mice) mice. eYFP vs. TeNT: *P* = 0.0917 (*t_4_* = 2.209). **f.** Time spent in shelter of n eYFP- (n = 3 mice) and TeNT- (n = 3 mice) mice after the onset of noise representation. eYFP vs. TeNT: *P* = 0.0411 (*t_4_* = 2.97). ICx, the cortex of the inferior colliculus. Unpaired *t* test (two-tailed) was applied in (d, e, f) for statistical analysis when data was normally distributed. Data are represented as mean (lines or bars) ± SEM (shaded regions or error bars); \**P* < 0.05, \*\**P* < 0.01. See also Supplementary Table 1.

## Discussion

In present study, we report that VGluT2+ neurons in the ATm, which is part of the non-lemniscal thalamic system, demonstrated sleep-wake state-dependent auditory (tone) or visual (blue light) responses. This is distinct from the largely preserved sensory responses observed in the lemniscal thalamic nuclei. Photoactivation of ATm^VGluT2+^ neurons induced immediate sleep-to-wake transition and pupil dilation in sleeping and awake behaving mice, respectively. Both ATm^VGluT2+^ neurons and their projections to the TeA/ECT were indispensable for the arousal induced by sensor stimuli of both modalities, while ATm^VGluT2+^ projections to tested subcortical structures showed modality preference in terms of arousing effect. Moreover, ATm^VGluT2+^ excitation in freely-moving mice induced behaviors signaling anxiety. Blocking ATm^VGluT2+^ synaptic transmission selectively and completely abolished sound-evoked escape behavior without affecting exploratory behavior. Our findings strongly suggest that the ATm is an interface between alerting sensory stimuli and arousal associated with defensive emotional state, which is required for the initiation of sensory-evoked innate defensive behaviors.

### The lemniscal thalamocortical pathway is not involved in arousal induction

Photoactivating the ATm induced immediate NREM sleep-to-wake transition, while stimulating the MGBv or dLGN did not. This distinction is very interesting because it has long been assumed that the lemniscal thalamic nuclei and their cortical projections are able to promote sensory evoked arousal. We thought this unexpected result could be explained by the distinct projection patterns of these two parallel thalamic systems and related structures.

In terms of cortical projection, ATm^VGluT2+^ neurons mainly project to the TeA/ECT ^21^ and, interestingly, predominantly target pyramidal neurons, especially those located in L5 (Supplementary Fig. 4B). Thus, the projection neurons in the TeA/ECT could presumably be directly and strongly activated by the thalamic inputs arising from the ATm. Since TeA/ECT^Pyr^ neurons project to widespread cortical areas including the prefrontal cortices ^27^, they possess a potential to produce global cortical activation. More than that, TeA/ECT^Pyr^ neurons also make substantial projections to subcortical regions critically involved in arousal and consciousness control, such as the nucleus accumbens ^45, 46, 47^ and claustrum ^48, 49^ (Supplementary Fig. 4D). Regarding subcortical projection, a high density of ATm^VGluT2+^ axonal fibers were observed in but not limited to the VMH and TS ^21^, which are important for emotional and behavioral state regulation. Our photoinhibition data suggested that the activity in ATm^VGluT2+^-VMH and -TS pathways contributed to sensory evoked arousal as well. Collectively, ATm^VGluT2+^ neurons can promote arousal via multiple parallel downstream pathways.

The projection pattern of the lemniscal thalamus such as the MGBv is much simpler than that of the ATm. Lemniscal thalamic neurons almost only project to the primary sensory cortices besides their innervation of the thalamic reticular nucleus. Sensory information from the lemniscal thalamus arrives in layer 4 (L4) first ^55^ and then propagate through the cortical column along the L4→L2/3→L5/6 pathway. In the auditory system, although L5 neurons in the Au1 make direct projections to the TeA/ECT ^42^, their axonal fibers preferentially target the superficial layer rather than the middle or deep layers (Allen Mouse Brain Atlas, connectivity. brain-map.org/projection/experiment/146858006), which receive strong inputs from ATm^VGluT2+^ neurons. Therefore, it is very possible that the group of TeA/ECT neurons receiving Au1 inputs are distinct from the TeA/ECT^pyr^ neurons that are innervated by ATm^VGluT2+^ neurons, thereby not being involved in arousal induction. In addition to corticocortical projections, Au1 L5 neurons also project to subcortical structures such as the ICx and TS ^56, 57^, which sends outputs to and receives inputs from the ATm ^21^, respectively. Further neural mechanistic dissection at the local circuit level may be needed to shed light on why the MGBv-Au1-ICx or -TS pathways fail to promote arousal.

### Relationship between the ATm and other wakefulness-promoting thalamic nuclei

Disorders of consciousness have long been observed in patients that suffered from direct injuries to their central thalamus ^58, 59, 60^, suggesting a critical role of central thalamus in maintaining general arousal. Recently published studies provided causal evidences for this proposal and revealed underlying mechanisms ^18, 20, 47^. The ATm might be able to activate certain subregions of the central thalamus by means of indirect projections, thereby facilitating global thalamic and cortical activation. For example, we demonstrated that the ATm made dense projections to the VMH, and that these projections had a significant contribution to blue light-induced sleep-to-wake transition. Since it has been reported that the VMH can indirectly activate the NAc via the midline PVT ^61^ and the PVT-NAc pathway can promote arousal ^47^, the ATm-VMH-PVT-NAc pathway might be one of the effector pathways contributing to blue light-induced arousal. Whether the ATm also contributes to the control of general arousal remains an open question.

### Relationship between the ATm and ascending reticular activating system

Our results showed that although photoinhibition of ATm^VGluT2+^ neurons greatly reduced the probability of sensory evoked awakening, it did not completely block sleep-to-wake transitions, suggesting the involvement of other pathways. Indeed, the level of tonic activity of noradrenergic neurons in the locus coeruleus (LC) can regulate the threshold of sensory evoked awakening ^62^, and the magnitude of sensory stimuli-evoked responses of dopaminergic neurons in the dorsal raphe nucleus (DRN) is significantly correlated with the probability of sensory evoked awakening ^63^. Both the LC and DRN are members of the ARAS in the brainstem, and theoretically, they are capable of promoting arousal via both ascending and descending pathways ^1, 3, 64, 65^. How is the ATm connected with the LC and DRN then? A recent retrograde tracing study shows that although the projection arising from the LC is negligible, the input from the DRN is noticeable ^21^. However, due to the existence of multiple cell types including DA neurons in the DRN, it is still unclear which type or types of DRN cells project to the ATm and how these projections may contribute to ATm-mediated sensory evoked arousal.

### The role of ATm^VGluT2+^ neurons in sound-triggered escape behavior

Selectively blocking the synaptic transmission of ATm^VGluT2+^ neurons, which receive inputs from the ICx ^21^, abolished escape behavior in response to suddenly presented noise without affecting locomotion (Fig. 6c, d). A previous study reported that the ICx is essential for sound-triggered running in head-fixed mice ^57^, which mimics sound-triggered flight behavior in freely-moving preparation. However, different from our findings, this study suggested that the projections from the ICx to the dorsal periaqueductal gray (dPAG) is the major downstream pathway. In addition, two other studies have shown that projections from the auditory cortex and deep layers of the SC to the dPAG, respectively, also contribute significantly to sound-evoked escape behavior ^66, 67^. Our results suggest that although there exist multiple brain regions, including the ICx, relaying alerting auditory information to the dPAG, the arousal and defensive emotional state promoted by ATm^VGluT2+^ neurons may strongly modulate the excitability of dPAG neurons in awake behaving mice, which are required for the initiation of defensive behaviors ^68^. Being presumably gated by ATm^VGluT2+^ neurons under normal conditions, excessive activation of dPAG neurons could be avoided, thus preventing behavioral hypersensitivity to environmental auditory stimuli or even stimuli of other modalities. It would be of great interest to test this hypothesis experimentally in the future because it may bear clinical significance for autistic children, a large fraction of whom are behaviorally characterized by sensory hypersensitivity associated with hyperarousal ^69^.

## Acknowledgments

We thank Z. X. Liu for immumohistochemical staining support, and W. J. Cheng for behavioral experiment assistance, W. Xiong and B. Hong for instrumental support, M. M. Luo for VGluT2-Cre mice and viruses supplement, and Z. C. Guo, H. L. Hu, Z. L. Huang, Y. Wang, Z. J. Xiao and Y. Q. Yu for critical comments on the manuscript. We also would like to thank Beijing Innovation Center for Future Chip at Tsinghua University for their technical support. K. Y. receives funding from National Natural Science Foundation of China (31871057, 81527901), Beijing Municipal Science & Technology Commission (Z181100001518004, Z181100001518006), and Guoqiang Institute, Tsinghua University. B.H. is supported by National Key R&D Program of China (2017YFA0205904).

## Author contributions

K.Y. conceptualized the study. Y.W., L.Y., K.T. designed parts of the study and conducted a majority of the experiments including related data collection and analysis. M.L. assisted with various recordings and analyzed data with W.H.. J.Z. assisted with optogenetic manipulation experiments. T.L. and R.Y. conducted sound-induced defensive behavioral experiments and related data analysis. F.X. and F.X. conducted head-fixed behavioral experiments and related data analysis. Y.G. assisted with anatomical experiments. Z.Y. and C.S. under the supervision of P.C. assisted with behavioral experiments. All authors contributed to data interpretation. K.Y. supervised the work together with B.H., and wrote the paper with Y.W., L.Y., K.T., M.L..

## Methods

### Mice

All animal procedures were approved by the Institutional Animal Care and Use Committee (IACUC) of Tsinghua University, Beijing, China. VGluT2-IRES-Cre mice obtaining from The Jackson Laboratory (Stock# 028863) were backcrossed to C57BL/6J wild-type to generate experimental heterozygous mice (abbreviated as VGluT2-IRES-Cre for convenience). Both male wild-type (WT) and heterozygous VGluT2-IRES-Cre mice (2–4 months of age) used in this study were housed under a 12h:12h light-dark cycle (light on from 10 a.m. to 10 p.m.) with ad libitum access to food and water.

### Virus preparation

All recombinant adeno-associated virus (AAV) vectors of 2/9 serotype (AAV2/9-hSyn-DIO-GCaMP6m, AAV2/9-hEF1α-DIO-hChR2(H134R)-EYFP, AAV2/9-CAG-DIO-taCaspase3, AAV2/9-CAG-EBFP2, and AAV2/9-hEF1a-DIO-mCherry), 2/5 serotype (AAV2/5-hSyn-hChR2(H134R)-mCherry, AAV2/5-hSyn-EGFP), 2/8 serotype (AAV2/8-hEF1α-GCaMP6m) and scAAV2/1-hSyn-Cre were purchased from Shanghai Taitool Bioscience Co. Ltd. (Shanghai, China). Cre-dependent AAV2/9 serotype encoding GtACR1 (AAV2/9-CAG-DIO-GtACR1-GFP) was kindly gifted by Minmin Luo Lab, and AAV2/9-CMV-DIO-GFP-2A-TeNT was packaged from Peng Cao Lab. In addition, AAV1-CBA-Flex-Arch-GFP, AAV1-hEF1α-DIO-eYFP, AAV1-CAG-Flex-eGFP, AAV1-hSyn-GCaMP6m were purchased from the University of Pennsylvania Vector Core (UPENN; Philadelphia, USA). AAV1-CAG-GFP was bought from University of North Carolina Vector Core (UNC; Carpel Hill, USA), and rAAV-hSyn-NES-jRGECO1a was bought from Wuhan BrainVTA Co. Ltd. (Wuhan, China). The final viral vector titers were diluted to a range of 3-5E +12 V.G/ml.

### General surgical procedures and viral delivery

Mice were first anesthetized with sodium pentobarbital (50 mg/kg, intraperitoneal injection, i.p.) and placed to a stereotaxic apparatus (RWD, 68001, Shenzhen, China). Standard animal surgery was strictly performed to expose brain surface above the ATm, MGBv or dLGN. Here, the coordinates used for ATm injection were anteroposterior (AP) −3.20 mm, mediolateral (ML): ±1.85 mm, dorsoventral (DV): −3.68 mm, where coordinates (AP/ML/DV) for MGBv and dLGN injection were −3.20 mm/−2.1 mm/−3.2 mm and −2.46 mm/−2.2 mm/−2.8 mm. All AAV vectors were stereotaxically injected with a glass pipette (4878; World Precision Instruments (WPI), Sarasota, FL, USA) by using a micro-syringe pump (UMP3; WPI, Sarasota, FL, USA) via micropipette (NanoFil; WPI, Sarasota, FL, USA) connected to a controller (Micro4; WPI, Sarasota, FL, USA). To avoid potential backflow, the pipette was slowly withdrawn at least 6 minutes after viral injection. After completion of any surgical procedures, mice were allowed to regain consciousness on a heating pad, and were transferred to a new individual residence room for full recovery (2-4 weeks) prior to the beginning of any behavioral operations.

For fiber photometry recording, the ATm of VGluT2-IRES-Cre mice was injected with AAV2/9-hSyn-DIO-GCaMP6m (200 nl), whereas the MGBv of WT mice were injected with either AAV1-Syn-GCaMP6m (150 nl) or AAV2/8-hEF1α-GCaMP6m (150 nl). Since it is difficult to define the borders distinctly between PIN and MGBv, we decided to insert the optical fiber ventromedially to PIN, and dorsolaterally to MGBv (similar coordinates to the virus injection site), which are coordinately different based on the Allen Mouse Brain Atlas. Moreover, the fiber tip was placed 0.25 mm above each injection site, to ensure that the normal auditory responses (by primary MGBv neurons) are preserved, although other non-lemniscal thalamic regions (MGBd/MGBm/MGBv for PIN; SG for MGBv) might cause certain damages due to the fiber placement. Above surgical consideration assumed that the neuronal activity recording and modulation would be more precisely, and the possible impact of virus injection titers and light power on the surrounding non-target neurons would be reduced. To verify the efficacy of ChR2 activation in MGBv, AAV2/5-hSyn-hChR2(H134R)-mCherry was used to activate MGBv, while rAAV-hSyn-NES-jRGECO1a was injected to record the neuronal activities in the primary auditory cortex (Au1). The ceramic ferrule was secured to skull surface at Au1 with dental cement.

For photoactivation of cell body experiments, AAV2/9-hEF1α-DIO-hChR2(H134R)-EYFP (200 nl) was injected to the ATm of VGluT2-IRES-Cre mice, whereas AAV2/5-hSyn-hChR2(H134R)-mCherry (70 nl) was injected to either MGBv or dLGN of WT mice. For the photoinhibition of ATm neurons, AAV2/9-CAG-DIO-GtACR1-GFP (200 nl) was injected to unilateral side of ATm, whereas the contra side of ATm was intermixed with AAV2/9-CAG-DIO-taCaspase3 and AAV2/9-CAG-DIO-GtACR1-GFP (total volume injection: 200 nl), therefore resulting in cell apoptosis within the ATm-targeted injection. All optogenetic experiments were started 2-3 weeks after the surgical operation. Besides, VGluT2-IRES-Cre mice injected with AAV1-hEF1α-DIO-EYFP or AAV1-CAG-FLEX-eGFP viruses (100 nl) in ATm, and WT mice injected with AAV2/5-hSyn-EGFP (70 nl) in MGBv or dLGN were used as control variable group for optogenetic manipulation.

To activate axonal fibers of VGluT2+ neurons, Cre-dependent AAV2/9-hEF1α-DIO-hChR2(H134R)-EYFP (200 nl) was delivered to ATm. For photoinhibition of axonal fibers, AAV1-CBA-FLEX-Arch-GFP (200 nl) was injected into unilateral side of ATm, whereas AAV2/9-CAG-DIO-taCaspase3 was intermixed with AAV1-CBA-FLEX-Arch-GFP (total volume injection: 200 nl) and injected into the contra side of ATm. After viral injection, an optical fiber was positioned at the dura mater surface of the TeA/ECT (posterior 3.2 mm from Bregma, lateral 0.2 mm from the rhinal fissure). After the skull surface of the target position was thinned, the optical fiber was secured to the skull surface using a thin layer of common dental cement. In addition, a second layer of graphite-mixed dental cement was applied to firm the fiber and prevent light leakage from reaching the eyes. After that, all mice were inspected for their optical fiber trace, and those who were not located in targeted position were excluded. Moreover, an optical fiber was implanted unilaterally above the VMH (AP/ML/DV: −1.5 mm/−0.4 mm/−5.2 mm) and TS (AP/ML/DV: −1.70 mm/−3.4 mm/−3.4 mm). For photoactivation of ATm-projecting terminals, AAV2/9-hEF1α-DIO-hChR2(H134R)-EYFP (70 nl) was unilaterally injected to ICx (AP/ML/DV coordinates for dorsal part of IC: −4.96mm/−0.50mm/−1.50mm and external part of IC: −4.96mm/−1.60mm/−2.00mm) and SO/dSC (AP/ML/DV coordinates: −4.36 mm/−0.5 mm/−1.2 mm), followed by the optical fiber implantation at ATm.

To label the axon fibers emanate from TeA/ECT, AAV1-CAG-GFP (200 nl) was delivered to TeA/ECT of WT mouse. For transsynaptic tracing of ATm-TeA/ECT pathway, AAV-hSyn-Cre intermixed with AAV-CAG-EBFP2 (mixed volume: 70 nl) were injected to the ATm of WT mice. Subsequently, AAV2/9-hEF1α-DIO-mCherry (300 nl) was used to target TeA/ECT neurons that specifically receive inputs from ATm. To visualize the cell-type of TeA/ECT neurons, brain slices expressing mCherry signal were selected to co-stain with GABA-specific antibodies.

### EEG and EMG implantation

To implant EEG/EMG apparatus, two craniotomy holes were drilled at the frontal cortical area (AP/ML: +1.5mm/+1.5mm) and the parietal area (for reference and ground; AP/ML: +3.2mm/+2.8mm) of the right hemisphere, respectively. Two miniature stainless screws, which served as EEG electrodes, were carefully inserted into the skull above the cortex, and two EMG electrodes were placed within the left and right trapezius muscles respectively, to monitor muscular activity. All EEG/EMG apparatus were secured to the skull using dental cement.

### Fiber photometry

To record Ca^2+^ signals from neuronal population level of targeted area, laser beam initially directed from 488-nm laser (OBIS 488LS; Coherent) was reflected through a dichroic mirror (MD498; Thorlabs), focused by a X10 objective lens (NA = 0.3; Olympus) and coupled to an optical commutator (Doric Lenses). A patch cord (diameter: 200 μm, NA = 0.37, 3-meters long) was used to guide the laser beam between the commutator and the implanted optical fiber in mice. To minimize GCaMP bleaching effect, the laser power was first adjusted at the tip of optical fiber to a low level of 0.02–0.03 mW. Next, GCaMP-coupling Ca^2+^ signals were bandpass filtered (MF525-39, Thorlabs) and collected by a photomultiplier tube (R3896, Hamamatsu). Then, an amplifier (C7319, Hamamatsu) was used to convert current output from the photomultiplier tube to voltage signal, which was further filtered through a low-pass filter (40 Hz cut-off; Brownlee 440). The analog voltage signal was digitalized at 1000 Hz by Power 1401 digitizer and analyzed with Spike2 software (CED, Cambridge, UK).

To measure the neuronal activity changes accurately, photometry data recording Ca^2+^ signals of neuronal population level were exported to Mat files in Spike2 and were processed with custom-written codes in MATLAB. The photometry signal F was converted to ΔF/F = (F-F_1_)/(F_1_-F_0_), where F_0_ and F_1_ represent the values of GCaMP-coupling Ca^2+^ signals before and after patch cord connection to mice at the beginning of the experiment, respectively.

To examine whether the sensory evoked neuronal activity correlates with corresponding brain states, ΔF/F data consisting of relevant events (e.g., onset of neuronal responses to sensory stimuli event was identified by recognizing TTL pulses) were extracted. We divided the results of 4 kHz tone and blue light stimulation into AN (awakened from NREM sleep), UAN (un-awakened from NREM sleep), and Wake trials, and averaged the ΔF/F values across mice based on the three categories above. Furthermore, we statistically compared the averaged ΔF/F values between the baseline (5 seconds before the sensory stimulation onset) and sensory-triggered events, where the duration of sensory-triggered events was calculated based on the average latency of cortical activation corresponding to each sensory modality (5 seconds for 4 kHz tone and 10 seconds for blue light) respectively. All ΔF/F values were plotted as mean value, whereas shaded area indicating SEM.

### Auditory- and visual-induced awakening test

In order to reduce environmental interference, all behavior experiments were conducted in a sound-attenuating chamber (IAC-Acoustics, UK). The background noise of the chamber is between 35 and 40 dB.

To examine the neuronal activity in response to sensory stimuli across sleep-wake state, either pure tone (4 kHz; 65 dB SPL; 10 s duration) or blue light (460-470 nm, 800 lx, 30 s duration) was applied in a pseudo-random order with an inter-stimulus-interval (ISI) of 8-15 minutes (4 to 5 trials for each stimulus). The speaker and LED light were located 50 cm above the center of the operant box and both stimuli were triggered and calibrated by a TDT System 3 (Tucker-Davis Technologies, Alachua, USA). At the same time, TTL time-locked pulses in response to either 4 kHz tone or blue light stimuli were generated and inputted into a Power 1401 digitizer and Spike2 software (CED, Cambridge, UK) remotely for accurate time stamping.

To differentiate the neuronal responses that corresponds to each brain states, TTL pulses were extracted and sleep-wake states were classified by another experimenter (Meijie Li), who was blinded to the experimental conditions. According to the behavioral consequences of sensory stimuli, each trial was classified as AN or UAN. For AN trials, experimental mice should show at least 3 seconds of EEG desynchronization from the delta or theta rhythms, and is occasionally accompanied by a sudden elevation of muscle activity for EMG within the sensory-stimulation time window. If these criteria were not met, the trial was categorized as UAN.

### Optogenetic modulation of states

One side of a patch cord (1.25 mm ceramic ferrule, 200 μm diameters, Inper) was connected to the implanted optical fiber, and another side (FC/PC connector) was connected to a swivel commutator (fiber-optic rotary joints, Inper) to allow free movement of mice. Then, patch cord that connected to the commutator was further linked to a blue-light or yellow-light laser emitter (MLL-FN-473nm/150 mW, Changchun New Industries Optoelectronics Technology, Changchun, China), which was controlled by RZ6 Multi-I/O Processor (Tucker-Davis Technologies, Alachua, USA). The intensity of the laser beam was measured with an optical power meter (PM100D, Thorlabs). For the laser power measurement, 1 or 10 mW laser was emitted at fiber tip for cellular stimulation, whereas 5-30 mW laser power was delivered for axonal stimulation.

For photoactivation experiments, 20 seconds of 473 nm laser pulses were delivered to activate the targeted ChR2-expressing neurons. Each laser stimulus was applied pseudo-randomly at ISI of 10-15 minutes, and around 40 trials with EEG/EMG signals were collected.

The photoinhibition of sensory-induced awakening tests were conducted after mice had fully recovered from surgery. On the first day, only 4 kHz tone or blue light was given to mice during stable NREM sleep (at least 10 seconds of continuous NREM sleep; similar to the fiber photometry experiments), and the probability of mice being awakened was recorded. For the photoinhibition tests, a continuous laser (473 nm for GtACR1; 593 nm for Arch) was given for 1 minute during stable NREM sleep, which the operations were carried out two days later after the baseline tests. In particular, 10 seconds of 4 kHz tone or 30 seconds of blue light was given at 20 seconds after the laser stimulus started, and the probability of cortical and behavioral activation within the sensory-stimulation time window were recorded.

### EEG-EMG recordings and analysis

We used Cerebus™ System (Blackrock Microsystems, Salt Lake City, USA) to record EEG/EMG signals in photoactivation and fiber photometry experiments. For photoinhibition experiments, we used RM6240 series multi-channel physiological signal acquisition and analysis system (Chengdu Instrument Factory, Chengdu, China) to record EEG/EMG signals. All the EEG/EMG signals were digitized at a sampling rate of 1000 Hz and were bandpass-filtered (EEG: 0.5-49 Hz, EMG: 20-500 Hz). Data analysis was mainly performed by the investigator (Meijie Li) who was blinded to the experimental manipulations.

The brain states were classified using custom written algorithm in MATLAB (MathWorks) based on the frequency and amplitude characteristics of the EEG and EMG, and corrected manually, if required. Low-amplitude EEG with low delta power (0.5-4 Hz) associated with elevated phasic EMG activity was categorized as wake, and high-amplitude EEG with high delta power associated with relatively lessened EMG activity was categorized as NREM sleep. A state of low-amplitude EEG with scarcely any EMG activity was categorized as REM sleep if regular theta waves (6-9 Hz) were visibly displayed in those EEG power spectra and a high theta/delta ratio value was calculated. All states were classified across consecutive non-overlapping 4 seconds windows.

In addition, for optogenetic inhibition experiments, similar criteria were used in fiber photometry tests to identify the behavioral consequences of sensory stimuli (to be classified as AN and UAN). Since the sensory-evoked awakening is not absolutely accompanied by significant EMG activity, thus we included the criterion of at least 3 seconds of continuous low-amplitude EEG desynchronization without significant EMG activity as the definition of cortical activation. For behavioral activation, a definition of cortical activation coupled with an increase in muscle tone (mainly from the head-neck movement in EMG activity) within the stimuli time window was used ^35, 63^. Relative delay represents the time difference between behavioral activation and cortical activation.

Before sleep stage scoring, EEG signals were bandpass filtered (0.5-49 Hz) and EMG signals were high pass filtered (25 Hz) respectively using FIR (Finite Impulse Response) filter, which was designed by the windowing method. Initially, we used Short-Time Fourier transform (STFT) with windows of 4 seconds and half-window-size overlapping to get the EEG spectrogram ^70^. For each time bin, we extracted 4 main features. EEG-based features composed of delta power (0.5-4 Hz), theta/delta ratio (6-9 Hz total power/0.5-4 Hz total power) and maximum amplitude of absolute value, whereas EMG-based feature represented the variance of EMG in each time bin ^32^.

Each set of extracted features was scaled to have a mean value of zero and standard deviation of one. Following feature extraction, we utilized EEG maximum amplitude of absolute value, delta power and EMG variance to construct a three-dimensional feature vector and implemented *k*-means algorithm with correlation distance to classify the states into two categories: roughly NREM and roughly wake. Then we separated REM from the above two categories with the criterion following: segments of which theta/delta ratio was greater than one were classified as REM. With the method described above, we obtained all 4-seconds periods to be clearly assigned to one of the sleep-wake states. These classification results were corrected by a supplementary rule: REM-sleep periods between wake periods were identified as wake ^71^. All the digital filters were designed by FDATool (Signal Processing Toolbox) in MATLAB, *k*-means algorithm was performed with in-built functions of the Statistics and Machine Learning toolbox in MATLAB.

For the calculation of percentage in each state across the photoactivation time window, a total of 220 seconds (including 20 seconds of light stimulation) was extracted and categorized into defined states as stated above. The state probability of each time bin per mouse was calculated as the number of trials in each state (NREM/REM/Wake) divided by the number of all trials, and was represented in percentage form to visually show an proportional changes of each state across each time bin in one figure ^63^. To determine the effect of photoactivation of ATm^VGluT2+^ neurons during NREM and REM sleep, we calculated the cortical activation latency which was defined as the time difference between the onset of laser ON and the time point indicating cortical activation. Since the time resolution based on state classification is 4 seconds, thus we determine the time point of cortical activation based on visual and spectral characteristics of the EEG/EMG signals.

For spectral analysis in photoactivation experiments, we extracted stimulated events (100 s before and after the stimulation period) to analyze certain frequency band power during photoactivation. For each event, EEG was decomposed into a time-frequency plane using the STFT with 4-seconds window and 80% overlap with a frequency resolution of 0.25 Hz. EEG delta frequency band power for each epoch was extracted by the integration of corresponding power over delta frequency band.

Mean delta power of all the stimulated events per mouse was normalized using z-score. This normalization corrected the variations in recorded signals across mice. We statistically compared the averaged delta power between baseline and the segments during and after perturbation. Furthermore, to enhance visualization, power of delta frequency band was smoothed by a 15-point gaussian filter with standard deviation of two. Epochs containing high amplitude artifacts were excluded from spectral analysis.

## Behavioral assessments

### Anxiety test

Open field (OF, 45 cm × 45 cm × 45 cm) and elevated plus maze (EPM, 50 cm × 10 cm for closed arms, 50 cm for platform height) tests were utilized to measure the anxiety level in ATm-photoactivating mice. Mice were initially placed in the operant chamber for a 5-minutes of exploratory and adaptation phase (Pre-OFF). Next, laser pulses (473 nm, 1mW, 10 Hz, 10 ms) were delivered immediately after the Pre-OFF session for another 5 minutes (ON) in both OF and EPM. Mice were then recorded for 5 minutes of post-stimulation phase (Post-OFF). All movement traces of mice were recorded with a camera, which further analyzed with custom-written MATLAB code to quantify the total number of entries, time spent duration and average speed of each mouse.

### Real-time place avoidance (RTPA)

The apparatus (50 cm × 25 cm × 30 cm) was comprised of two identical sides (25 cm × 25 cm) that were connected by an opening passage (10 cm width) at the center. At the beginning, mice were placed in either one side (starting side was counterbalanced across mice) for 5 minutes. Then, for the following 5 minutes, laser pulses (473 nm, 1 mW, 10 Hz, 10 ms) were applied automatically when the mouse entered the laser stimulation zone, whereas the laser stimulation stopped after the mouse had left. Another post-session (Post-OFF) was given for 5 minutes to examine the behavioral patterns of the mouse. Similar to the assessments of OF and EPM, all movement traces were recorded with a camera, which further analyzed with custom-written MATLAB code to quantify the total number of entries, time spent duration and average speed of each mouse.

### Noise-induced escape behavior

To block the synaptic transmission of ATm^VGluT2+^ neurons receiving from the ICx, scAAV2/1-hSyn-Cre was bilaterally injected in the ICx, and later AAV2/9-CMV-DIO-GFP-2A-TeNT was injected in the ATm of both sides.

For noise-induced escape behavior tests, mice were initially placed in an open field (35 cm × 35 cm × 35 cm), in which a sheltered nest (20 cm × 12 cm × 15 cm) was provided at the corner of the chamber ^56^. When mice were exploring in the center zone, a white noise (75 dB SPL, 10 s) was delivered from a speaker located above the chamber ^72^. Generally, mice exhibited escape behavior to white noise immediately and concealed itself inside the shelter for the rest of stimulation time. All movement traces were recorded with two cameras from two adjacent sides, which further analyzed offline with custom-written MATLAB code to calculate the speed of each mouse throughout the noise-triggered escape events. To compare the distinct behavioral response of TeNT and eYFP mice, the peak (highest) speed of each mouse was extracted within defined time phase: Pre (−2∼0 s), Noise (0∼2 s), Noise (2∼10 s). Besides, latency to escape and time spent in shelter for each mouse were analyzed and calculated as well.

### Pupillometry detection and data analysis

To perform the pupil detection in an awake mouse, a head plate was attached at the skull using dental cement after the virus injection and optical fiber implantation, which aligned with the coronal suture.

After 2-3 weeks of recovery, mouse head was fixed, and was obligated to acclimate on a cylindrical treadmill. Meanwhile, the right eye of mouse was detected by an infrared camera, and the RZ6 Multi-I/O Processor (Tucker-Davis Technologies, Alachua, USA) was used to manipulate the photoactivation moment to the head-fixed mouse by controlling the blue-light laser emitter (MLL-FN-473 nm/150 mW, Changchun New Industries Optoelectronics Technology, Changchun, China) with a patch cord connecting to the optical fiber of mouse. During the photoactivation event, laser pulses (similar to the photoactivation of ATm^VGluT2+^ neurons) were delivered to the ATm^VGluT2+^ neurons for 5 seconds, and every ISI was given for 25 seconds.

Time-stamp of pupil frames aligned with the photoactivation-triggered events were extracted and analyzed with custom-written MATLAB code. Pupil frames were first extracted at 20 Hz, and were used to detect the pupil position by comparing with the pre-defined pixel threshold. The pupil diameter (pixels) was determined as the major axis of the ellipse using ellipsoid fitting algorithm. To quantify the ATm photoactivating effects on pupil dilation, we compared the pupil diameters of mice at two different time point (the photoactivation onset vs. the photoactivation offset). All 10 trials of pupil diameters per mouse were normalized to the range of [0, 1], and every normalized diameter value of each mouse at defined time point were averaged, thus to display the tendency of pupil dilation induced by the photoactivation of ATm^VGluT2+^ neurons.

### Histology and Immunohistochemistry

Mice were deeply anaesthetized with pentobarbital sodium and were transcardially perfused with 0.9% saline followed by 4% paraformaldehyde. For fixation, brains were kept overnight in 4% paraformaldehyde. For cryoprotection, brains were placed in 30% sucrose (w/v) in PBS solution for at least two nights. After embedding and freezing, the mouse brain was sectioned coronally (35 mm thick) with a cryostat (Leica CM1900). For Nissl staining, we followed the instruction of the Nissl Staining kit (Beyotime Biotechnology, Shanghai, China). For immunofluorescent staining, brain slices were first incubated with PBST solution (PBS containing 0.3 % of Triton X-100) for 5 minutes 3 times each to permeabilize cell membranes. Then, the slices were rinsed with PBS solution for 3 times, followed by a blocking buffer (10 % donkey serum in PBST solution) for 90 minutes at room temperature. Subsequently, the blocking buffer was replaced with diluted anti-GABA (1:1000, rabbit, Sigma-Aldrich; diluted with blocking buffer), and the slices were kept at 4℃ overnight. After rinsing 3 times with PBS, the slices were incubated with the Alexa 488-conjugated secondary antibodies (1:500, goat, Abcam) for 2 hours at room temperature. Once again, the slices were rinsed with PBS for 3 times and 15 minutes each. Lastly, all slices were transferred onto a glass side with coverslip mounted on top. At this stage, the slides were scanned to obtain the images of the targeted fluorescence signals.

### Statistical analysis

All data were presented as mean ± SEM. Sample sizes were determined based on the previous publications that used optogenetics to study sleep-wake state regulation ^63^. All statistical analysis were performed with MATLAB (Mathworks) or Prism8 (GraphPad). We first performed the Shapiro-Wilk test on each dataset for normality test. Parametric tests were used if the dataset pass the normality test. Otherwise, non-parametric tests were used, especially in sensory-induced awakening experiments (due to the variance of individual differences to sensory response or limitation of collecting trials). For data of two groups, we used paired two-tailed Student’s *t* tests if the data was normally distributed, and unpaired two-tailed Student’s *t* test if the data was normally distributed and had homogeneity of variance. If two groups of data were not normally distributed, Mann-Whitney *U* test or Wilcoxon signed rank test was used for statistical analysis. Data was considered to be statistically significant if it passed \**P* < 0.05, \*\**P* < 0.01, \*\*\**P* < 0.001 and \*\*\*\**P* < 0.0001.

---

**Supplementary Fig. 1.**
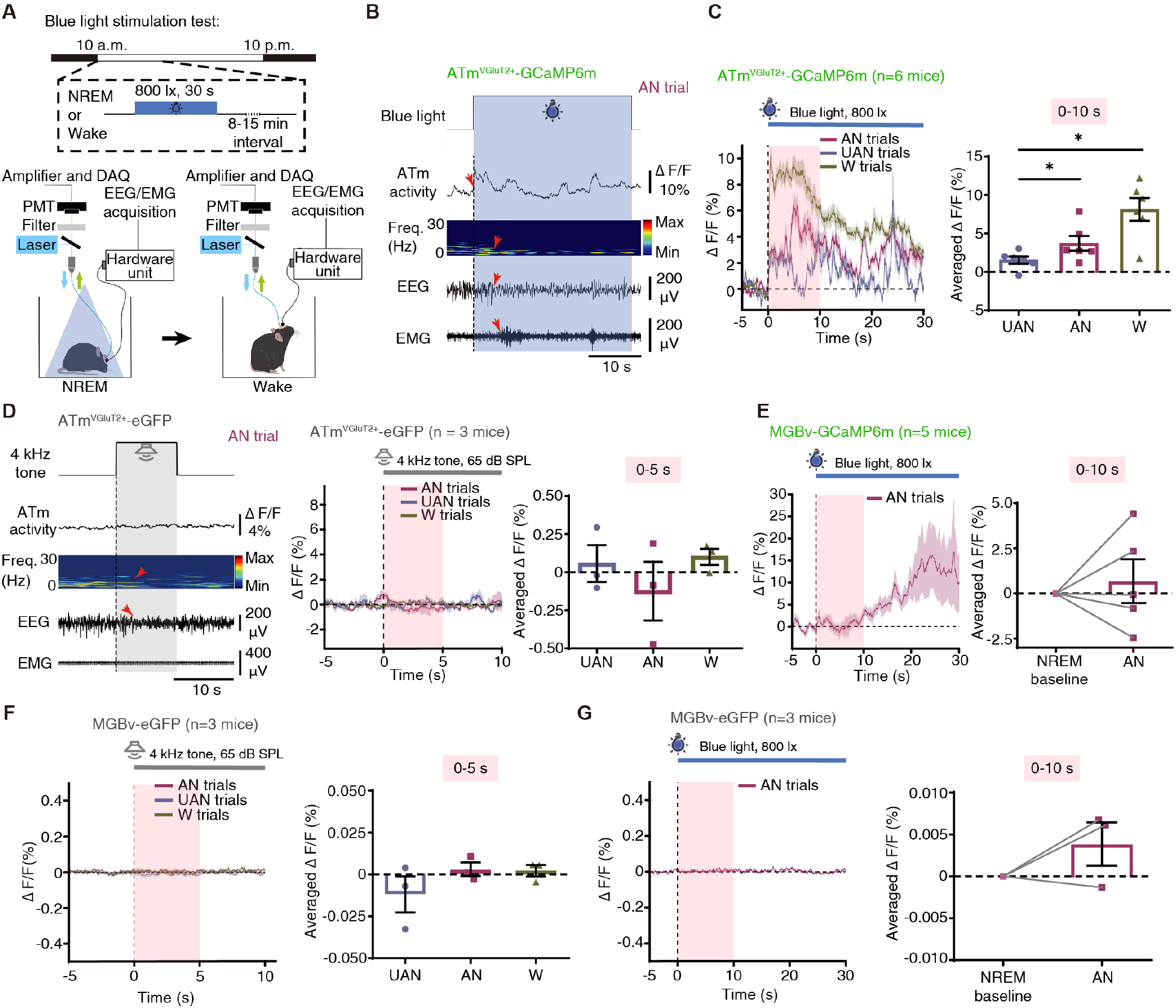
Related to Fig. 1. Sensory responsiveness of ATmVGluT2+ but not MGBv neuronal population correlates with sleep-wake states. **(A)** Upper panel: Schematic diagram illustrating the parametric protocol for presenting blue light stimulation. A blue light (800 lx, 30 s duration) was manually applied to mice with inter-stimulus-interval (ISI) of 8-15 minutes during light cycle (10:00 a.m.-10:00 p.m.). Bottom panel: Schematic design showing simultaneous monitoring of ATm^VGluT2+^ activity induced by blue light and recording of EEG/EMG signals in a freely-moving mouse across sleep-wake states. **(B)** Blue light-induced ATm^VGluT2+^ activity, EEG power spectrum, and EEG/EMG signals in an example awakened (AN) trial. Vertical dotted line indicates the onset of blue light stimulation; Translucent blue area indicates the duration of blue light. Red arrows from top to bottom indicate the onset of blue light-induced ATm^VGluT2+^ activity, delta (0.5-4 Hz) power reduction, cortical activation, and behavioral activation, respectively. Scale bar: 10 seconds. **(C)** Left: Averaged ATm^VGluT2+^ activity in AN, un-awakened (UAN) and wakefulness (W) trials. Vertical dotted line indicates the onset of stimulation; Translucent pink area indicates the time period (10 seconds) within which most blue light-induced awakenings occurred. Right: Averaged ΔF/F values in the first 10 seconds after blue light onset in three trial types. *n* = 6 mice, UAN vs. AN: *P* = 0.0313 (*W* = 21); UAN vs. W: *P* = 0.0152 (*U* = 3). **(D)** Left: Tone-induced EEG power spectrum, and EEG/EMG signals in an example AN trial of eGFP mouse. Scale bar: 5 seconds. Middle: Averaged ATm^VGluT2+^ activity in AN, UAN and W trials of eGFP mice. Right: Averaged ΔF/F values in the first 5 seconds after tone onset in three trial types. **(E)** Left: Averaged MGBv activity in AN trial. Right: Averaged ΔF/F values of 5 seconds before and 10 seconds after the blue light onset in AN trial. **(F)** Left: Averaged MGBv activity in AN, UAN and W trials of eGFP mice. Right: Averaged ΔF/F values in the first 5 seconds after tone onset in three trial types. **(G)** Left: Averaged MGBv activity in AN trial of eGFP mice. Right: Averaged ΔF/F values of 5 seconds before and 10 seconds after the blue light onset in AN trial. Paired *t* test (two-tailed) was used in (e, f, g) when data are normally distributed, and Wilcoxon matched-pairs signed rank test or Mann-Whitney *U* test was applied in (c, d, e, f) for statistical analysis. Data are represented as mean (lines or bars) ± SEM (shaded regions or error bars); \**P* < 0.05. See also Supplementary Table 1.

**Supplementary Fig. 2.**
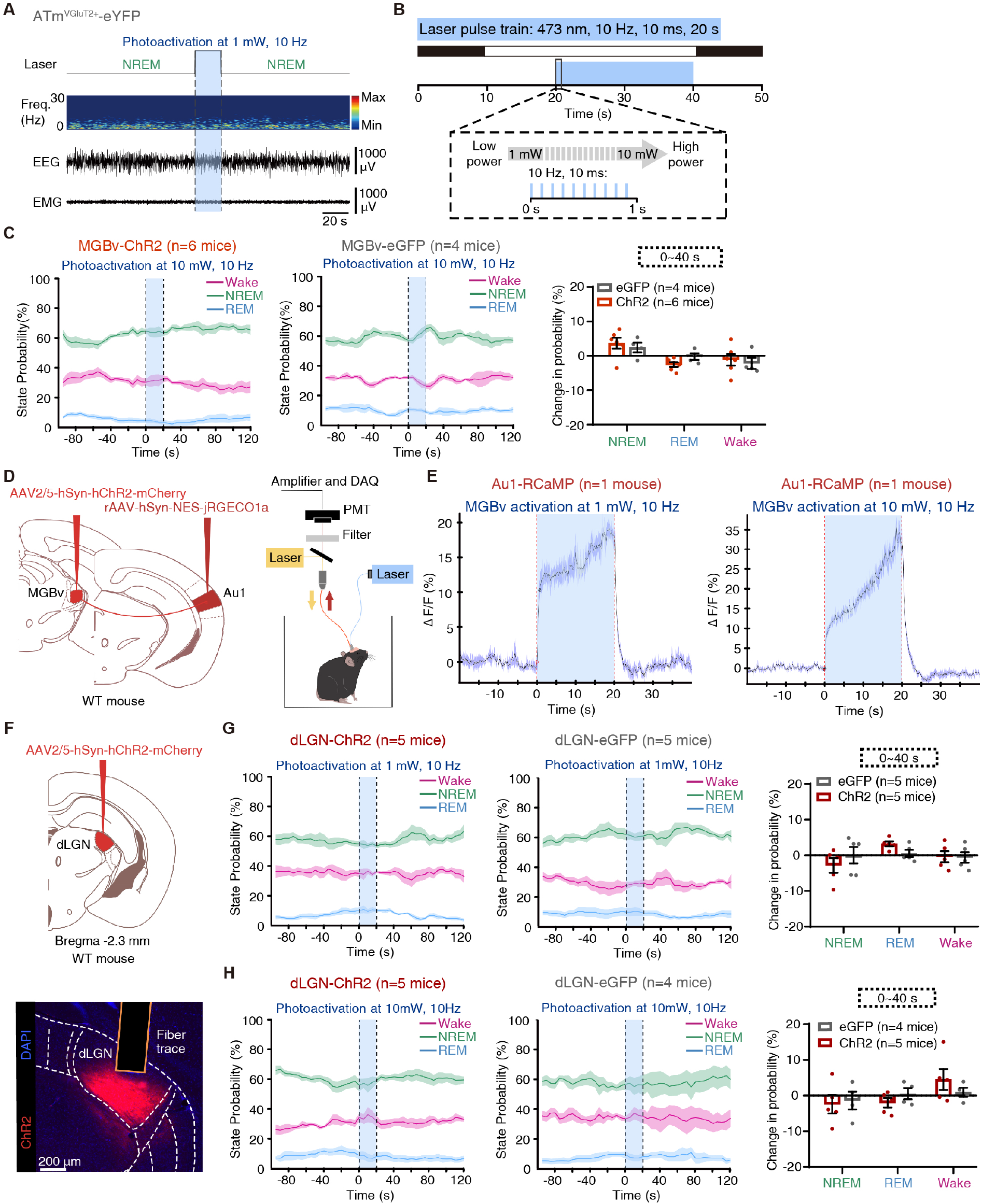
Expanded data to Fig. 2. Photoactivation of the MGBv or dLGN does not induce NREM sleep-to-wake transition. **(A)** EEG power spectrum and EEG/EMG signals in a representative trial showing un-awakened effect after photoactivation of ATm^VGluT2+^ neurons in eYFP mouse. **(B)** Schematic diagram illustrating the parametric protocol for photoactivation. Laser pulses (473 nm, 10 Hz, 10 ms, 20 s duration) were delivered at either 1 mW or 10 mW during light cycle (10:00 a.m.-10:00 p.m.). **(C)** State probability before, during and after MGBv photoactivation in ChR2- (left, *n* = 6 mice) and eGFP- (middle, *n* = 4 mice) mice during light cycle (10:00 a.m.-10:00 p.m.). Right: Averaged changes in state probability within 40 seconds after photoactivation onset. **(D)** Schematic design showing simultaneous photoactivation of MGBv neurons and recording of ipsilateral Au1 activity in a freely-moving mouse. **(E)** Averaged Au1 activity in response to low-power (left) and high-power (right) activation of MGBv neurons. **(F)** Upper panel: Schematic of virus injection in the dLGN of WT mouse for photoactivation. Lower panel: Representative expression of ChR2 and fiber trace for photoactivation in the dLGN. Scale bar: 200 μm. **(G-H)** State probability before, during and after low-power **(G)** and high-power **(H)** photoactivation of dLGN in ChR2- (left, *n* = 5 mice) and eGFP- (middle, *n* = 5 or 4 mice) mice during light cycle. Right: Averaged changes in state probability within 40 seconds after photoactivation onset. dLGN, the dorsal lateral geniculate nucleus. Unpaired *t* test (two-tailed) was used in (c, g, h), otherwise Mann-Whitney *U* test was applied for statistical analysis. Data are represented as mean (lines or bars) ± SEM (shaded regions or error bars). See also Supplementary Table 1.

**Supplementary Fig. 3.**
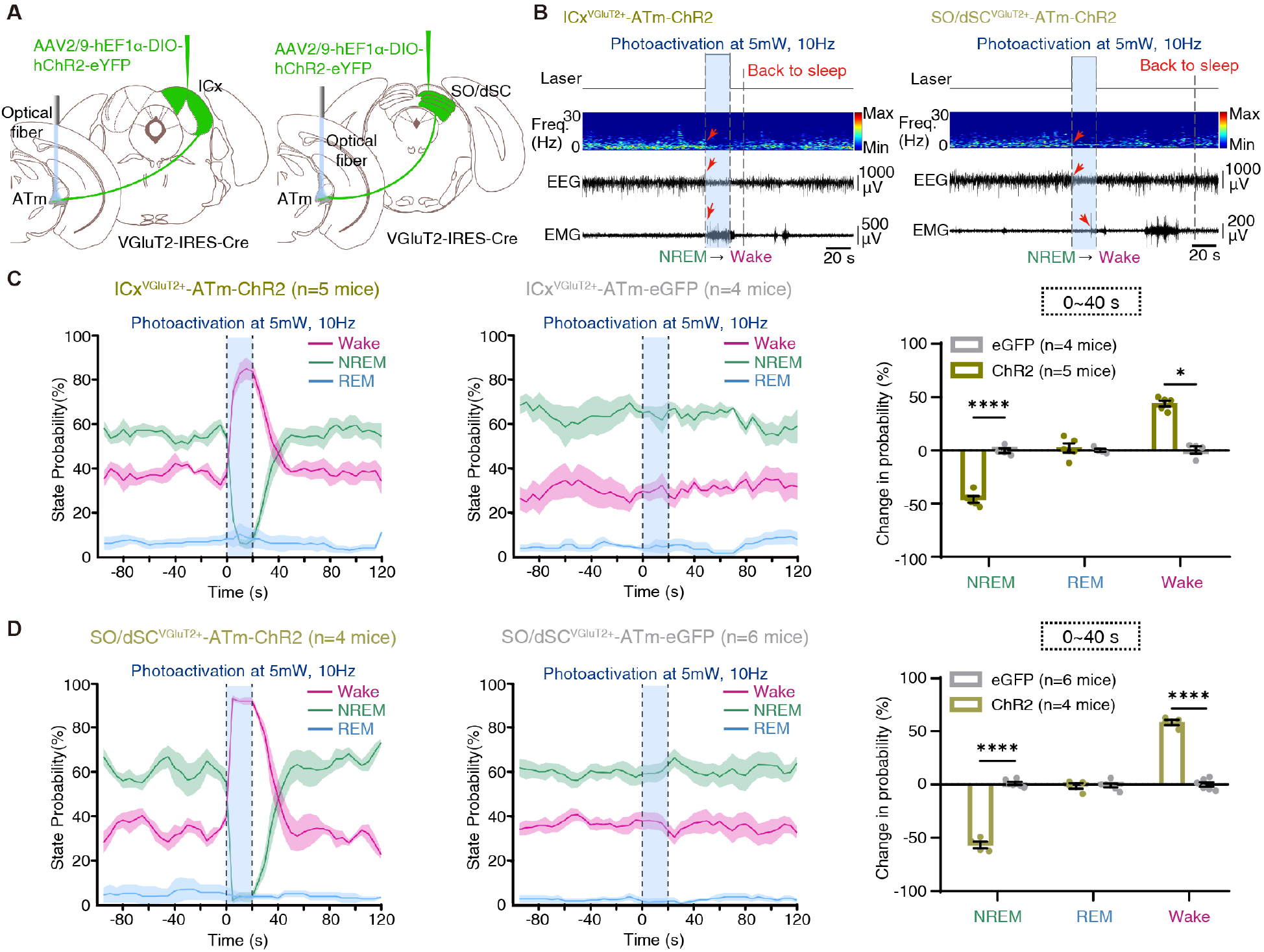
Expanded data to Fig. 3. Photoactivation of the ICx^VGluT2+^- or SO/dSC^VGluT2+^-ATm pathways induce rapid NREM sleep-to-wake transition. **(A)** Schematic of experimental set-up for photoactivation of ICx^VGluT2+^ (left) and SO/dSC^VGluT2+^ (right) axonal fibers in the ATm. **(B)** EEG power spectrum and EEG/EMG signals in a representative trial showing ICx^VGluT2+^ (left) and SO/dSC^VGluT2+^ (right) photoactivation-induced NREM-to-Wake transition. Translucent light-blue area indicates the duration of photoactivation; Red arrows from top to bottom indicate the onset of delta power reduction, cortical activation, and behavioral activation, respectively. Scale bar: 20 seconds. **(C)** State probability before, during and after ICx^VGluT2+^ photoactivation in ChR2- (left, *n* = 5 mice) and eGFP- (middle, *n* = 4 mice) mice during light cycle. Right: Averaged changes in state probability within 40 seconds after photoactivation onset. NREM sleep, *P* < 0.0001 (*t_7_* = 11.32); Wake, *P* = 0.0159 (*U* = 0). **(D)** State probability before, during and after SO/dSC^VGluT2+^ activation in ChR2- (left, *n* = 4 mice) and eGFP- (middle, *n* = 6 mice) mice during light cycle. Right: Averaged changes in state probability within 40 seconds after photoactivation onset. NREM sleep, *P* < 0.0001 (*t_8_* = 18.96); Wake, *P* < 0.0001 (*t_8_* = 17.59). ICx, the cortex of the inferior colliculus; SO/dSC, the optic and deeper layers of the superior colliculus. Unpaired *t* test (two-tailed) was used in (c, d) when data are normally distributed, otherwise Mann-Whitney *U* test was applied for statistical analysis. Data are represented as mean (lines or bars) ± SEM (shaded regions or error bars); \**P* < 0.05, \*\*\*\**P* < 0.0001. See also Supplementary Table 1.

**Supplementary Fig. 4.**
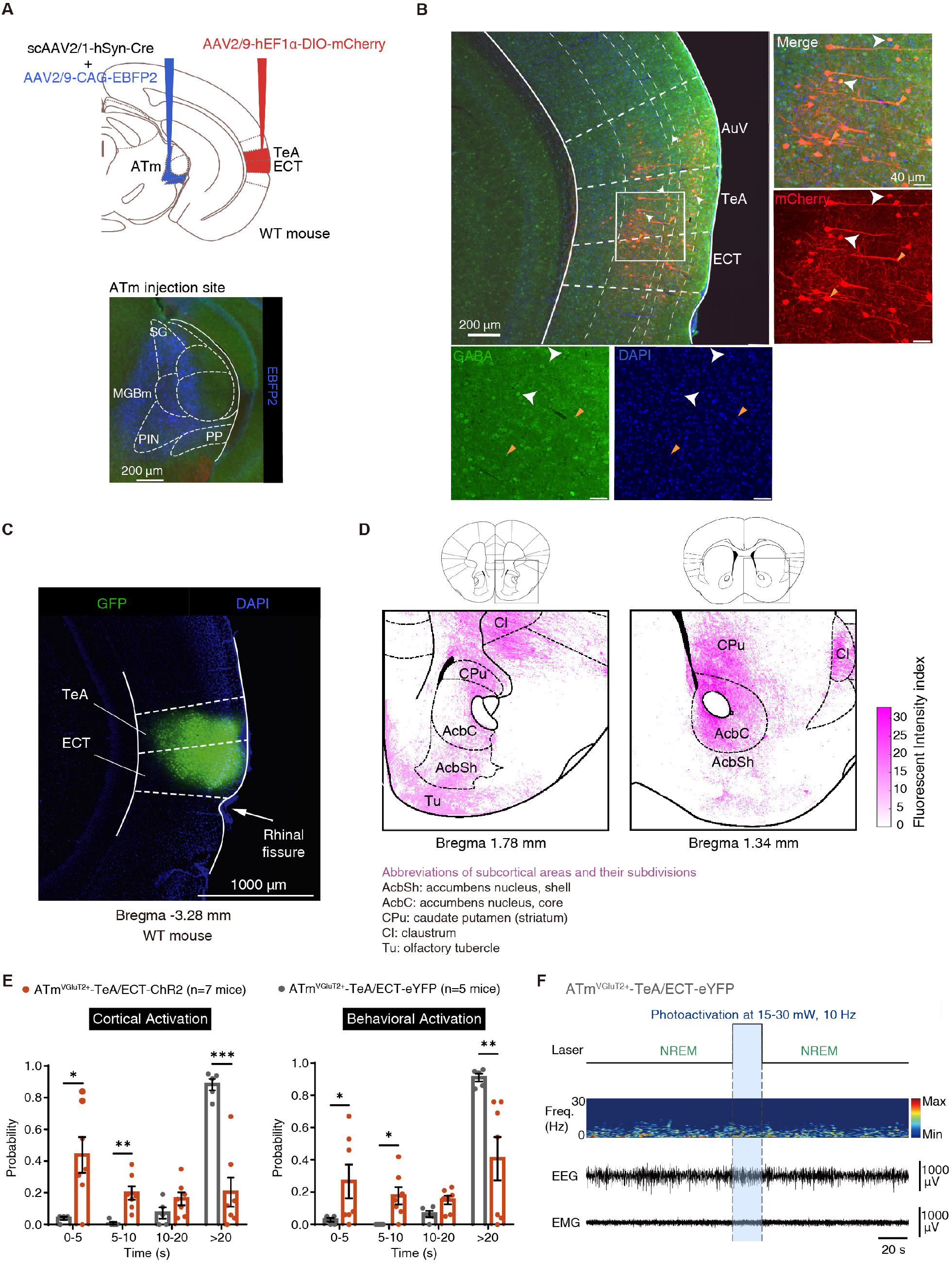
Expanded data to Fig. 4. Transsynaptic labeling of and the downstream target of TeA/ECT neurons that may participate in cortical and behavioral activation of ATmVGluT2+-TeA/ECT pathway excitation. **(A)** Left: Diagram showing virus injection strategy enabling anterograde trans-monosynaptic expression of mCherry in TeA/ECT neurons receiving ATm inputs. Right: Expression of EBFP2 in the ATm. Scale bar: 200 μm. **(B)** Left: Representative field of the cortex showing expression of mCherry and GABA. DAPI marked all nuclei in the field. Right: Magnified view of the boxed area in the left panel, showing TeA/ECT neurons that express mCherry. Scale bar: 200 μm (left) or 40 μm (right). White arrows indicate mCherry and GABA colocalized neurons; yellow arrows indicate neurons that express mCherry only. **(C)** Expression of GFP in TeA/ECT neurons. **(D)** Monosynaptic outputs in cortical and subcortical regions of TeA/ECT in a WT mouse. **(E)** Probability of cortical (left) and behavioral (right) activation at four consecutive time period after ATm^VGluT2**+**^ axonal photoactivation during NREM sleep in ChR2- (Orange, *n* = 7 mice) and eYFP- (Gray, *n* = 5 mice) mice. Left: 0-5 s, *P* = 0.029 (*U* = 4.5); 5-10 s, *P* = 0.0025 (*U* = 0); >20 s, *P* = 0.0001 (*t_10_* = 5.915). Right: 0-5 s, *P* = 0.0316 (*U* = 4); 5-10 s, *P* = 0.0114 (*U* = 2.5); >20 s, *P* = 0.0025 (*U* = 0). **(F)** EEG power spectrum and EEG/EMG signals in a representative trial showing un-awakened effect after photoactivation of ATm^VGluT2+^ axonal fibers in eYFP mouse. ATm, the medial sector of the auditory thalamus; PIN, the posterior intralaminar thalamic nuclei; PP, the peripeduncular nucleus; MGBm, the medial division of the medial geniculate body; SG, the suprageniculate nucleus; AuV, the ventral area of secondary auditory cortex; TeA, temporal association cortex; ECT, ectorhinal cortex. Unpaired *t* test (two-tailed) was used in (a) when data are normally distributed, otherwise Mann-Whitney *U* test was applied for statistical analysis. Data are represented as mean (bars) ± SEM (error bars); \**P* < 0.05, \*\**P* < 0.01, \*\*\**P* < 0.001. See also Supplementary Table 1.

**Supplementary Fig. 5.**
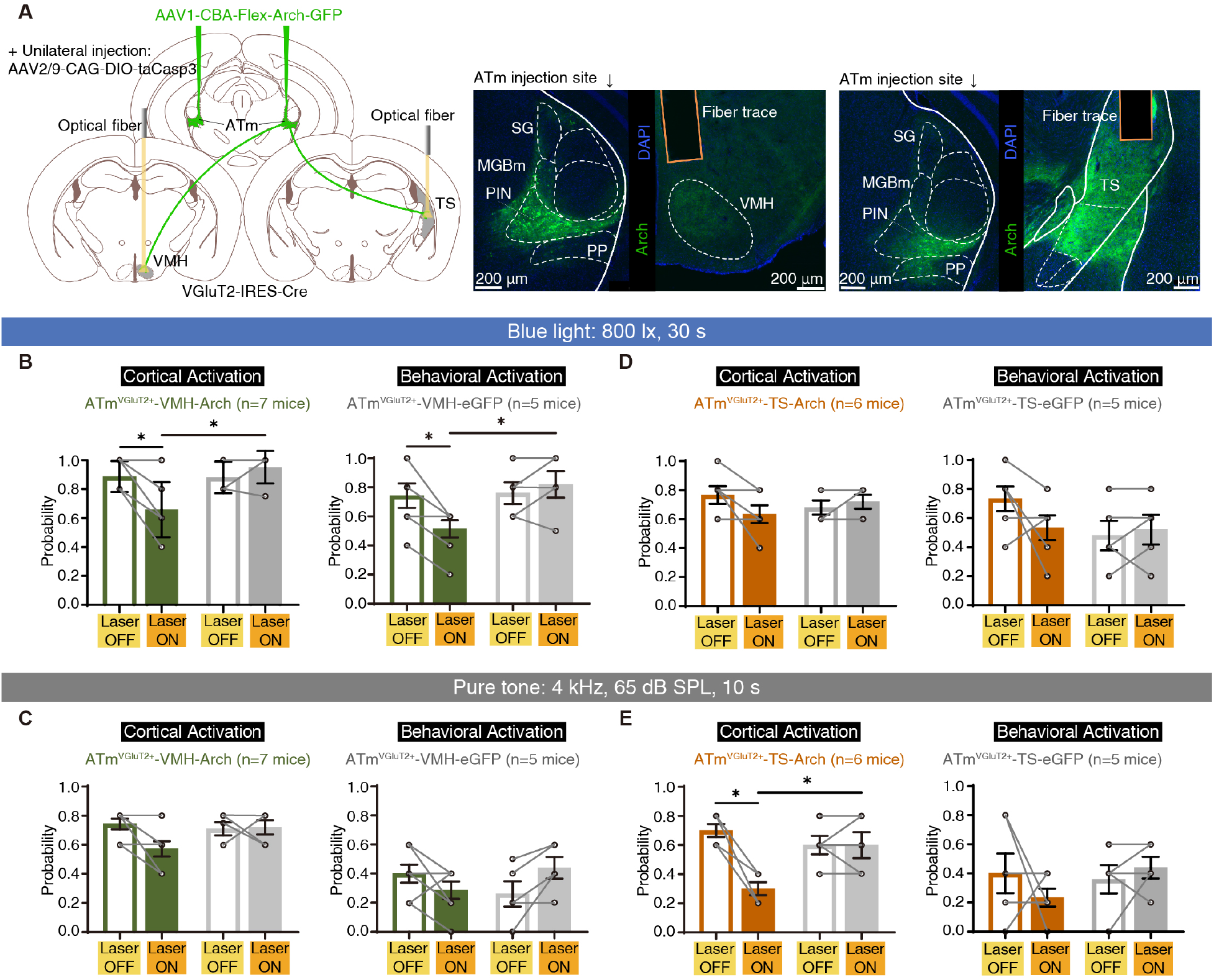
Expanded data to Fig. 4. The contribution of ATm^VGluT2+^-VMH and - TS pathways to sensory evoked awakening shows modality preference. **(A)** Left: Schematic of experimental set-up for photoinhibition of ATm^VGluT2+^ axonal fibers in the VMH and TS. Right two panels: Representative expression of Arch in the ATm and fiber trace for photoinhibition in VMH (middle) or TS (right). **(B)** Probability of blue light-induced cortical (left) and behavioral (right) activation in Arch- (Olive green, *n* = 7 mice) and eGFP- (Gray, *n* = 5 mice) mice in Laser OFF and ON trials. Left: Arch, Laser OFF vs. Laser ON, *P* = 0.0313 (*W* = −21); Laser ON, Arch vs. eGFP, *P* = 0.0253 (*U* = 4). Right: Arch, Laser OFF vs. Laser ON, *P* = 0.0313 (*W* = −21); Laser ON, Arch vs. eGFP, *P* = 0.0202 (*U* = 5). **(C)** Probability of tone-induced cortical (left) and behavioral (right) activation in Arch- and eGFP-mice in Laser OFF and ON trials. **(D)** Probability of blue light-induced cortical (left) and behavioral (right) activation in Arch- (Dark orange, *n* = 6 mice) and eGFP- (Gray, *n* = 5 mice) mice in Laser OFF and ON trials. **(E)** Probability of tone-induced cortical (left) and behavioral (right) activation in Arch- and eGFP-mice in Laser OFF and ON trials. Left: Arch, Laser OFF vs. Laser ON, *P* = 0.0313 (*W* = −21); Laser ON, Arch vs. eGFP, *P* = 0.0433 (*U* = 3). VMH, the ventromedial hypothalamus; TS, the tail of the striatum. Paired or unpaired *t* test (two-tailed) was used in (b, c, d, e) when data are normally distributed, otherwise Wilcoxon matched-pairs signed rank test or Mann-Whitney *U* test was applied for statistical analysis. Data are represented as mean (bars) ± SEM (error bars); \**P* < 0.05. See also Supplementary Table 1.

**Supplementary Fig. 6.**
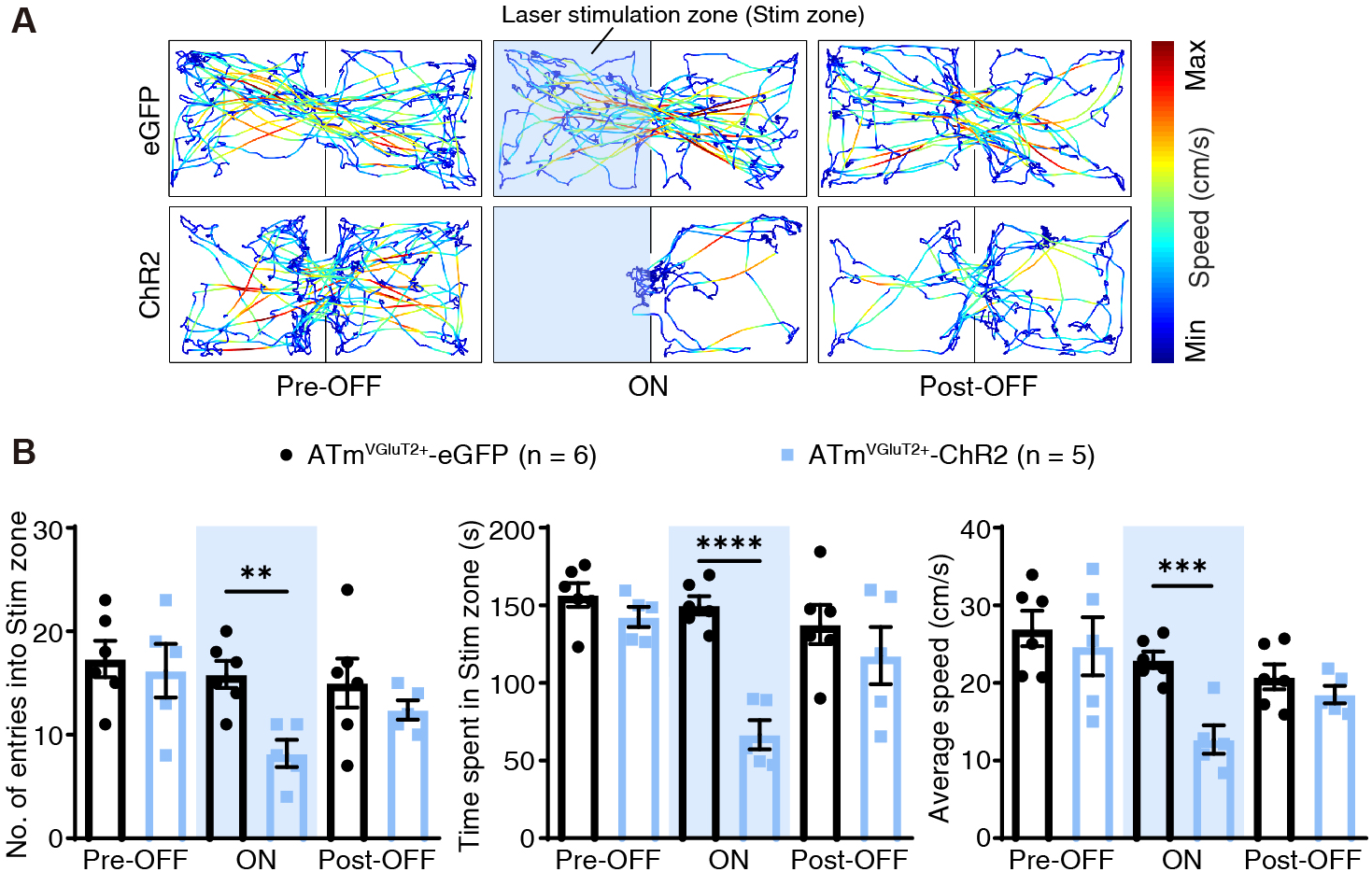
Related to Fig. 5. Photoactivation of ATm^VGluT2+^ neurons induces avoidance behavior in RTPA test. **(A)** Representative movement traces of eGFP- (upper row) and ChR2- (lower row) mice in a schematic real-time place avoidance (RTPA) before (left column, Pre-OFF), during (middle column, ON) and after (right column, Post-OFF) photoactivation. Translucent light blue area marks the laser stimulation zone (Stim zone). The color gradient indicates the magnitude of movement speed in RTPA test. **(B)** The number of entries into the Stim zone (left), time spent in the Stim zone (middle), and average speed (right) of ChR2- (Sky blue, *n* = 5 mice) and eGFP- (Black, *n* = 6 mice) mice. eGFP (ON) vs. ChR2 (ON), left to right, *P* = 0.0027 (*t_9_* = 4.087); *P* < 0.0001 (*t_9_* = 7.799); *P* = 0.0007 (*t_9_* = 5.051). Unpaired *t* test (two-tailed) was applied in (b) when data was normally distributed. Data are represented as mean (bars) ± SEM (error bars); \*\**P* < 0.01, \*\*\**P* < 0.001, \*\*\*\**P* < 0.0001. See also Supplementary Table 1.

**Supplementary Table 1. Statistical analysis relates to Fig. 1-6 and Supplementary Fig. 1-6.**

**Supplementary Movie 1. Photoactivation of ATm^VGluT2+^ neurons in ChR2 and eYFP mice.**

Low-power activation of ATm^VGluT2+^ neurons in ChR2 mouse (left) induces immediate cortical activation followed by behavioral activation, while did not show any abovementioned effects in eYFP mouse (right).

**Supplementary Movie 2. Photoactivation of MGBv or dLGN neurons in ChR2 mouse.**

Both 1 mW and 10 mW power activation of MGBv or dLGN neurons did not show any arousal-related signals in ChR2 mouse.

**Supplementary Movie 3. Sensory-induced changes in EEG/EMG signals in baseline and photoinhibition tests of GtACR1 mouse.**

Photoinhibition (Laser ON) of ATm^VGluT2+^ neurons hampered both blue light- and tone-induced cortical activation and behavioral activation in GtACR1 mouse.

**Supplementary Movie 4. Photoactivation of ATm^VGluT2+^ axonal fibers in TeA/ECT of ChR2 and eYFP mice.**

Photoactivation of ATm^VGluT2+^ axonal fibers in TeA/ECT of ChR2 mouse (left) induces cortical activation followed by behavioral activation, while did not show any abovementioned effects in eYFP mouse (right).

## References

1. McCormick DA, Nestvogel DB, He BJ. Neuromodulation of Brain State and Behavior. Annu Rev Neurosci 43, 391–415 (2020).

2. Scammell TE, Arrigoni E, Lipton JO. Neural Circuitry of Wakefulness and Sleep. Neuron 93, 747–765 (2017).

3. Lee SH, Dan Y. Neuromodulation of brain states. Neuron 76, 209–222 (2012).

4. Liu D, Dan Y. A Motor Theory of Sleep-Wake Control: Arousal-Action Circuit. Annu Rev Neurosci 42, 27–46 (2019).

5. Brown RE, Basheer R, McKenna JT, Strecker RE, McCarley RW. Control of sleep and wakefulness. Physiol Rev 92, 1087–1187 (2012).

6. Hua R, et al. Calretinin Neurons in the Midline Thalamus Modulate Starvation-Induced Arousal. Curr Biol 28, 3948–3959 e3944 (2018).

7. Yamagata T, et al. The hypothalamic link between arousal and sleep homeostasis in mice. Proc Natl Acad Sci U S A 118, (2021).

8. Neckelmann D, Ursin R. Sleep stages and EEG power spectrum in relation to acoustical stimulus arousal threshold in the rat. Sleep 16, 467–477 (1993).

9. Oswald I, Taylor AM, Treisman M. Discriminative responses to stimulation during human sleep. Brain 83, 440–453 (1960).

10. Tye KM. Neural Circuit Motifs in Valence Processing. Neuron 100, 436–452 (2018).

11. Hwang K, Bertolero MA, Liu WB, D’Esposito M. The Human Thalamus Is an Integrative Hub for Functional Brain Networks. J Neurosci 37, 5594–5607 (2017).

12. Ahissar E, Sosnik R, Haidarliu S. Transformation from temporal to rate coding in a somatosensory thalamocortical pathway. Nature 406, 302–306 (2000).

13. Guillery RW, Sherman SM. Thalamic relay functions and their role in corticocortical communication: generalizations from the visual system. Neuron 33, 163–175 (2002).

14. Winer JA, Miller LM, Lee CC, Schreiner CE. Auditory thalamocortical transformation: structure and function. Trends Neurosci 28, 255–263 (2005).

15. Steriade M. The corticothalamic system in sleep. Front Biosci 8, d878–899 (2003).

16. Issa EB, Wang X. Sensory responses during sleep in primate primary and secondary auditory cortex. J Neurosci 28, 14467–14480 (2008).

17. Portas CM, Krakow K, Allen P, Josephs O, Armony JL, Frith CD. Auditory processing across the sleep-wake cycle: simultaneous EEG and fMRI monitoring in humans. Neuron 28, 991–999 (2000).

18. Gent TC, Bandarabadi M, Herrera CG, Adamantidis AR. Thalamic dual control of sleep and wakefulness. Nat Neurosci 21, 974–984 (2018).

19. Honjoh S, Sasai S, Schiereck SS, Nagai H, Tononi G, Cirelli C. Regulation of cortical activity and arousal by the matrix cells of the ventromedial thalamic nucleus. Nat Commun 9, 2100 (2018).

20. Matyas F, et al. A highly collateralized thalamic cell type with arousal-predicting activity serves as a key hub for graded state transitions in the forebrain. Nat Neurosci 21, 1551–1562 (2018).

21. Cai D, et al. Distinct Anatomical Connectivity Patterns Differentiate Subdivisions of the Nonlemniscal Auditory Thalamus in Mice. Cereb Cortex 29, 2437–2454 (2019).

22. Gharaei S, Honnuraiah S, Arabzadeh E, Stuart GJ. Superior colliculus modulates cortical coding of somatosensory information. Nat Commun 11, 1693 (2020).

23. Oh SW, et al. A mesoscale connectome of the mouse brain. Nature 508, 207–214 (2014).

24. Anderson LA, Christianson GB, Linden JF. Stimulus-specific adaptation occurs in the auditory thalamus. J Neurosci 29, 7359–7363 (2009).

25. Lee CC. Exploring functions for the non-lemniscal auditory thalamus. Front Neural Circuits 9, 69 (2015).

26. Lu E, Llano DA, Sherman SM. Different distributions of calbindin and calretinin immunostaining across the medial and dorsal divisions of the mouse medial geniculate body. Hear Res 257, 16–23 (2009).

27. Zingg B, et al. Neural networks of the mouse neocortex. Cell 156, 1096–1111 (2014).

28. Gunaydin LA, et al. Natural neural projection dynamics underlying social behavior. Cell 157, 1535–1551 (2014).

29. Rajasethupathy P, Ferenczi E, Deisseroth K. Targeting Neural Circuits. Cell 165, 524–534 (2016).

30. Urban DJ, Roth BL. DREADDs (designer receptors exclusively activated by designer drugs): chemogenetic tools with therapeutic utility. Annu Rev Pharmacol Toxicol 55, 399–417 (2015).

31. Bourgin P, Hubbard J. Alerting or Somnogenic Light: Pick Your Color. PLoS Biol 14, e2000111 (2016).

32. Kohtoh S, Taguchi, Y., Matsumoto, N., Wada, M., Huang, Z.-L., and Urade Y. Algorithm for sleep scoring in experimental animals based on fast Fourier transform power spectrum analysis of the electroencephalogram. Sleep and Biological Rhythms 6, 163–171 (2008).

33. Ito T, Bishop DC, Oliver DL. Expression of glutamate and inhibitory amino acid vesicular transporters in the rodent auditory brainstem. J Comp Neurol 519, 316–340 (2011).

34. Cruikshank SJ, Killackey HP, Metherate R. Parvalbumin and calbindin are differentially distributed within primary and secondary subregions of the mouse auditory forebrain. Neuroscience 105, 553–569 (2001).

35. Halasz P, Terzano M, Parrino L, Bodizs R. The nature of arousal in sleep. J Sleep Res 13, 1–23 (2004).

36. Govorunova EG, Sineshchekov OA, Janz R, Liu X, Spudich JL. NEUROSCIENCE. Natural light-gated anion channels: A family of microbial rhodopsins for advanced optogenetics. Science 349, 647–650 (2015).

37. Kim YS, et al. Crystal structure of the natural anion-conducting channelrhodopsin GtACR1. Nature 561, 343–348 (2018).

38. Yang CF, et al. Sexually dimorphic neurons in the ventromedial hypothalamus govern mating in both sexes and aggression in males. Cell 153, 896–909 (2013).

39. Wiegert JS, Mahn M, Prigge M, Printz Y, Yizhar O. Silencing Neurons: Tools, Applications, and Experimental Constraints. Neuron 95, 504–529 (2017).

40. Shang C, et al. BRAIN CIRCUITS. A parvalbumin-positive excitatory visual pathway to trigger fear responses in mice. Science 348, 1472–1477 (2015).

41. Shin Yim Y, et al. Reversing behavioural abnormalities in mice exposed to maternal inflammation. Nature 549, 482–487 (2017).

42. Tasaka GI, et al. The Temporal Association Cortex Plays a Key Role in Auditory-Driven Maternal Plasticity. Neuron 107, 566–579 e567 (2020).

43. Amaral DG, Schumann CM, Nordahl CW. Neuroanatomy of autism. Trends Neurosci 31, 137–145 (2008).

44. Schumann CM, et al. Longitudinal magnetic resonance imaging study of cortical development through early childhood in autism. J Neurosci 30, 4419–4427 (2010).

45. Eban-Rothschild A, Rothschild G, Giardino WJ, Jones JR, de Lecea L. VTA dopaminergic neurons regulate ethologically relevant sleep-wake behaviors. Nat Neurosci 19, 1356–1366 (2016).

46. Luo YJ, et al. Nucleus accumbens controls wakefulness by a subpopulation of neurons expressing dopamine D1 receptors. Nat Commun 9, 1576 (2018).

47. Ren S, et al. The paraventricular thalamus is a critical thalamic area for wakefulness. Science 362, 429–434 (2018).

48. Crick FC, Koch C. What is the function of the claustrum? Philos Trans R Soc Lond B Biol Sci 360, 1271–1279 (2005).

49. Goll Y, Atlan G, Citri A. Attention: the claustrum. Trends Neurosci 38, 486–495 (2015).

50. Lazarus M, Chen JF, Urade Y, Huang ZL. Role of the basal ganglia in the control of sleep and wakefulness. Curr Opin Neurobiol 23, 780–785 (2013).

51. Yamagata T, et al. The role of the hypothalamus in cortical arousal and sleep homeostasis. bioRxiv, 2020.2005.2019.104521 (2020).

52. Vinck M, Batista-Brito R, Knoblich U, Cardin JA. Arousal and locomotion make distinct contributions to cortical activity patterns and visual encoding. Neuron 86, 740–754 (2015).

53. Silva BA, Gross CT, Graff J. The neural circuits of innate fear: detection, integration, action, and memorization. Learn Mem 23, 544–555 (2016).

54. Sando R, et al. Assembly of Excitatory Synapses in the Absence of Glutamatergic Neurotransmission. Neuron 94, 312–321 e313 (2017).

55. Douglas RJ, Martin KA. Neuronal circuits of the neocortex. Annu Rev Neurosci 27, 419–451 (2004).

56. Li Z, et al. Corticostriatal control of defense behavior in mice induced by auditory looming cues. Nat Commun 12, 1040 (2021).

57. Xiong XR, et al. Auditory cortex controls sound-driven innate defense behaviour through corticofugal projections to inferior colliculus. Nat Commun 6, 7224 (2015).

58. Castaigne P, Lhermitte F, Buge A, Escourolle R, Hauw JJ, Lyon-Caen O. Paramedian thalamic and midbrain infarct: clinical and neuropathological study. Ann Neurol 10, 127–148 (1981).

59. Krolak-Salmon P, Croisile B, Houzard C, Setiey A, Girard-Madoux P, Vighetto A. Total recovery after bilateral paramedian thalamic infarct. Eur Neurol 44, 216–218 (2000).

60. Schiff ND. Central thalamic contributions to arousal regulation and neurological disorders of consciousness. Ann N Y Acad Sci 1129, 105–118 (2008).

61. Engelke DS, et al. A hypothalamic-thalamostriatal circuit that controls approach-avoidance conflict in rats. Nat Commun 12, 2517 (2021).

62. Hayat H, et al. Locus coeruleus norepinephrine activity mediates sensory-evoked awakenings from sleep. Sci Adv 6, eaaz4232 (2020).

63. Cho JR, et al. Dorsal Raphe Dopamine Neurons Modulate Arousal and Promote Wakefulness by Salient Stimuli. Neuron 94, 1205–1219 e1208 (2017).

64. Jones BE. Arousal and sleep circuits. Neuropsychopharmacology 45, 6–20 (2020).

65. Saper CB, Fuller PM. Wake-sleep circuitry: an overview. Curr Opin Neurobiol 44, 186–192 (2017).

66. Evans DA, Stempel AV, Vale R, Ruehle S, Lefler Y, Branco T. A synaptic threshold mechanism for computing escape decisions. Nature 558, 590–594 (2018).

67. Wang H, et al. Direct auditory cortical input to the lateral periaqueductal gray controls sound-driven defensive behavior. PLoS Biol 17, e3000417 (2019).

68. Canteras NS. The medial hypothalamic defensive system: hodological organization and functional implications. Pharmacol Biochem Behav 71, 481–491 (2002).

69. Marco EJ, Hinkley LB, Hill SS, Nagarajan SS. Sensory processing in autism: a review of neurophysiologic findings. Pediatr Res 69, 48R–54R (2011).

70. Putze F, Muhl C, Lotte F, Fairclough S, Herff C. Editorial: Detection and Estimation of Working Memory States and Cognitive Functions Based on Neurophysiological Measures. Front Hum Neurosci 12, 440 (2018).

71. Costa-Miserachs D, Portell-Cortes I, Torras-Garcia M, Morgado-Bernal I. Automated sleep staging in rat with a standard spreadsheet. J Neurosci Methods 130, 93–101 (2003).

72. Zhang GW, Shen L, Zhong W, Xiong Y, Zhang LI, Tao HW. Transforming Sensory Cues into Aversive Emotion via Septal-Habenular Pathway. Neuron 99, 1016–1028 e1015 (2018).

