## Supplementary Table 1 for "A non-lemniscal thalamic interface connecting alerting sensory cues to internal states in mice"

### Supplementary Table 1. Statistical analysis relates to Fig. 1-6 and Supplementary Fig. 1-6.

| Figure | *n* number | | Normality (Shapiro-Wilk test) | Equal variance test | Statistic method | *P* value | Statistics value |
|---|---|---|---|---|---|---|---|
| Fig. 1e (right, UAN vs AN) | | ATm$^{VGluT2+}$-GCaMP6m mice (*n* = | Passed | | Paired *t* test (two-tailed) | *P* = 0.0209 | $t_6$ = 3.108 |
| Fig. 1e (right, UAN vs W) | | ATm$^{VGluT2+}$-GCaMP6m mice (*n* = | Passed | Passed | Unpaired *t* test (two-tailed) | *P* = 0.0065 | $t_{12}$ = 3.285 |
| Fig. 1e (right, AN vs W) | | ATm$^{VGluT2+}$-GCaMP6m mice (*n* = | Passed | Passed | Unpaired *t* test (two-tailed) | *P* = 0.4569 | $t_{12}$ = 0.7687 |
| Fig. 1h (right, UAN vs AN) | | MGBv-GCaMP6m mice (*n* = | Passed | | Paired *t* test (two-tailed) | *P* = 0.7890 | $t_4$ = 0.2860 |
| Fig. 1h (right, UAN vs W) | | MGBv-GCaMP6m mice (*n* = | Passed | Passed | Unpaired *t* test (two-tailed) | *P* = 0.5990 | $t_8$ = 0.5475 |
| Fig. 1h (right, AN vs W) | | MGBv-GCaMP6m mice (*n* = | Passed | Passed | Unpaired *t* test (two-tailed) | *P* = 0.5001 | $t_8$ = 0.7062 |
| Fig. 2c (right, OFF:-100-0 s) | ATm$^{VGluT2+}$-eYFP mice (*n* = 4) | ATm$^{VGluT2+}$-ChR2 mice (*n* = 7) | Failed | | Mann-Whitney *U* test (two-tailed) | *P* = 0.1727 | *U* = 6 |
| Fig. 2c (right, ON: 0-20 s) | ATm$^{VGluT2+}$-eYFP mice (*n* = 4) | ATm$^{VGluT2+}$-ChR2 mice (*n* = 7) | Passed | Passed | Unpaired *t* test (two-tailed) | *P* < 0.0001 | $t_9$ = 7.074 |
| Fig. 2c (right, OFF: 20-40 s) | ATm$^{VGluT2+}$-eYFP mice (*n* = 4) | ATm$^{VGluT2+}$-ChR2 mice (*n* = 7) | Passed | Passed | Unpaired *t* test (two-tailed) | *P* < 0.0001 | $t_9$ = 13.45 |
| Fig. 2c (right, OFF: 40-120 s) | ATm$^{VGluT2+}$-eYFP mice (*n* = 4) | ATm$^{VGluT2+}$-ChR2 mice (*n* = 7) | Passed | Passed | Unpaired *t* test (two-tailed) | *P* = 0.5527 | $t_9$ = 0.6168 |
| Fig. 2d (right, NREM) | ATm$^{VGluT2+}$-eYFP mice (*n* = 4) | ATm$^{VGluT2+}$-ChR2 mice (*n* = 7) | Passed | Passed | Unpaired *t* test (two-tailed) | *P* < 0.0001 | $t_9$ = 12.4 |
| Fig. 2d (right, REM) | ATm$^{VGluT2+}$-eYFP mice (*n* = 4) | ATm$^{VGluT2+}$-ChR2 mice (*n* = 7) | Passed | Passed | Unpaired *t* test (two-tailed) | *P* = 0.0022 | $t_9$ = 4.239 |
| Fig. 2d (right, Wake) | ATm$^{VGluT2+}$-eYFP mice (*n* = 4) | ATm$^{VGluT2+}$-ChR2 mice (*n* = 7) | Passed | Passed | Unpaired *t* test (two-tailed) | *P* < 0.0001 | $t_9$ = 13.93 |
| Fig. 2f (right) | ATm$^{VGluT2+}$-eYFP mice (*n* = 4) | ATm$^{VGluT2+}$-ChR2 mice (*n* = 10) | Passed | Passed | Unpaired *t* test (two-tailed) | *P* < 0.0001 | $t_{12}$ = 9.908 |
| Fig. 2g (NREM →Wake vs REM→Wake) | | ATm$^{VGluT2+}$-ChR2 mice (*n* = 7) | Passed | | Paired *t* test (two-tailed) | *P* = 0.0311 | $t_6$ = 2.803 |
| Fig. 2j (right, NREM) | MGBv-eGFP mice (*n* = 5) | MGBv-ChR2 mice (*n* = 6) | Failed | | Mann-Whitney *U* test (two-tailed) | *P* = 0.5368 | *U* = 11 |
| Fig. 2j (right, REM) | MGBv-eGFP mice (*n* = 5) | MGBv-ChR2 mice (*n* = 6) | Passed | Passed | Unpaired *t* test (two-tailed) | *P* = 0.8992 | $t_9$ = 0.1303 |
| Fig. 2j (right, Wake) | MGBv-eGFP mice (*n* = 5) | MGBv-ChR2 mice (*n* = 6) | Passed | Passed | Unpaired *t* test (two-tailed) | *P* = 0.8213 | $t_9$ = 0.2326 |
| Fig. 3e (left, Laser OFF vs Laser ON) | | ATm$^{VGluT2+}$-GtACR1 mice (*n* = 7) | Failed | | Wilcoxon matched-pairs signed rank test (two-tailed) | *P* = 0.0156 | *W* = -28 |
| | ATm$^{VGluT2+}$-eYFP mice (*n* = 5) | | Failed | | Wilcoxon matched-pairs signed rank test (two-tailed) | *P* > 0.9999 | *W* = 2 |
| Fig. 3e (left, eYFP vs GtACR1) | ATm$^{VGluT2+}$-eYFP mice (*n* = 5) | ATm$^{VGluT2+}$-GtACR1 mice (*n* = 7) | Laser OFF | | | | |
| | | | Failed | | Mann-Whitney *U* test (two-tailed) | *P* > 0.9999 | *U* = 15.5 |
| | | | Laser ON | | | | |
| | | | Failed | | Mann-Whitney *U* test (two-tailed) | *P* = 0.0051 | *U* = 0.5 |

| | | | | | | | |
|---|---|---|---|---|---|---|---|
| Fig. 3e (right, Laser OFF vs Laser ON) | | ATm$^{VGluT2+}$-GtACR1 mice (*n* = | Passed | | Paired *t* test (two-tailed) | *P* = 0.0249 | $t_{6}$ = 2.970 |
| | ATm$^{VGluT2+}$-eYFP mice (*n* = 5) | | Failed | | Wilcoxon matched-pairs signed rank test (two-tailed) | *P* = 0.8125 | *W* = 3 |
| Fig. 3e (right, eYFP vs GtACR1) | ATm$^{VGluT2+}$-eYFP mice (*n* = 5) | ATm$^{VGluT2+}$-GtACR1 mice (*n* = 7) | Laser OFF | | | | |
| | | | Failed | | Mann-Whitney *U* test (two-tailed) | *P* = 0.6136 | *U* = 14 |
| | | | Laser ON | | | | |
| | | | Passed | Passed | Unpaired *t* test (two-tailed) | *P* = 0.0115 | $t_{10}$ = 3.086 |
| Fig. 3f (left, Laser OFF vs Laser ON) | | ATm$^{VGluT2+}$-GtACR1 mice (*n* = 12) | Failed | | Wilcoxon matched-pairs signed rank test (two-tailed) | *P* = 0.0005 | *W* = -78 |
| | ATm$^{VGluT2+}$-eYFP mice (*n* = 9) | | Passed | | Paired *t* test (two-tailed) | *P* = 0.1791 | $t_{8}$ = 1.473 |
| Fig. 3f (left, eYFP vs GtACR1) | ATm$^{VGluT2+}$-eYFP mice (*n* = 9) | ATm$^{VGluT2+}$-GtACR1 mice (*n* = 12) | Laser OFF | | | | |
| | | | Failed | | Mann-Whitney *U* test (two-tailed) | *P* = 0.4037 | *U* = 42 |
| | | | Laser ON | | | | |
| | | | Passed | Passed | Unpaired *t* test (two-tailed) | *P* = 0.0015 | $t_{19}$ = 3.715 |
| Fig. 3f (right, Laser OFF vs Laser ON) | | ATm$^{VGluT2+}$-GtACR1 mice (*n* = 12) | Failed | | Wilcoxon matched-pairs signed rank test (two-tailed) | *P* = 0.0391 | *W* = -31 |
| | ATm$^{VGluT2+}$-eYFP mice (*n* = 9) | | Passed | | Paired *t* test (two-tailed) | *P* = 0.1776 | $t_{8}$ = 1.478 |
| Fig. 3f (right, eYFP vs GtACR1) | ATm$^{VGluT2+}$-eYFP mice (*n* = 9) | ATm$^{VGluT2+}$-GtACR1 mice (*n* = 12) | Laser OFF | | | | |
| | | | Passed | Passed | Unpaired *t* test (two-tailed) | *P* = 0.5420 | $t_{19}$ = 0.6209 |
| | | | Laser ON | | | | |
| | | | Failed | | Mann-Whitney *U* test (two-tailed) | *P* = 0.0412 | *U* = 26 |
| Fig. 4c (right, OFF -100-0 s) | ATm$^{VGluT2+}$-TeA/ECT-eYFP mice (*n* = 5) | ATm$^{VGluT2+}$-TeA/ECT-ChR2 mice (*n* = 7) | Passed | Passed | Unpaired *t* test (two-tailed) | *P* = 0.7088 | $t_{10}$ = 0.3844 |
| Fig. 4c (right, ON: 0-20 s) | ATm$^{VGluT2+}$-TeA/ECT-eYFP mice (*n* = 5) | ATm$^{VGluT2+}$-TeA/ECT-ChR2 mice (*n* = 7) | Passed | Passed | Unpaired *t* test (two-tailed) | *P* = 0.0022 | $t_{10}$ = 4.085 |
| Fig. 4c (right, OFF: 20-40 s) | ATm$^{VGluT2+}$-TeA/ECT-eYFP mice (*n* = 5) | ATm$^{VGluT2+}$-TeA/ECT-ChR2 mice (*n* = 7) | Passed | Passed | Unpaired *t* test (two-tailed) | *P* < 0.0001 | $t_{10}$ = 6.722 |
| Fig. 4c (right, OFF: 40-120 s) | ATm$^{VGluT2+}$-TeA/ECT-eYFP mice (*n* = 5) | ATm$^{VGluT2+}$-TeA/ECT-ChR2 mice (*n* = 7) | Passed | Passed | Unpaired *t* test (two-tailed) | *P* = 0.0683 | $t_{10}$ = 2.043 |
| Fig. 4d (right, NREM) | ATm$^{VGluT2+}$-TeA/ECT-eYFP mice (*n* = 5) | ATm$^{VGluT2+}$-TeA/ECT-ChR2 mice (*n* = 7) | Passed | Passed | Unpaired *t* test (two-tailed) | *P* = 0.0023 | $t_{10}$ = 4.069 |
| Fig. 4d (right, REM) | ATm$^{VGluT2+}$-TeA/ECT-eYFP mice (*n* = 5) | ATm$^{VGluT2+}$-TeA/ECT-ChR2 mice (*n* = 7) | Passed | Passed | Unpaired *t* test (two-tailed) | *P* = 0.1206 | $t_{10}$ = 1.697 |
| Fig. 4d (right, Wake) | ATm$^{VGluT2+}$-TeA/ECT-eYFP mice (*n* = 5) | ATm$^{VGluT2+}$-TeA/ECT-ChR2 mice (*n* = 7) | Passed | Passed | Unpaired *t* test (two-tailed) | *P* = 0.0004 | $t_{10}$ = 5.209 |

| | | | | | | | |
|---|---|---|---|---|---|---|---|
| Fig. 4i (left, Laser OFF vs Laser ON) | | ATm$^{VGluT2+}$-TeA/ECT-Arch mice ($n$ = 7) | Failed | | Wilcoxon matched-pairs signed rank test (two-tailed) | $P$ = 0.0313 | $W$ = -21 |
| | ATm$^{VGluT2+}$-TeA/ECT-eYFP mice ($n$ = 7) | | Failed | | Wilcoxon matched-pairs signed rank test (two-tailed) | $P > 0.9999$ | $W$ = 1 |
| Fig. 4i (left, eYFP vs Arch) | ATm$^{VGluT2+}$-TeA/ECT-eYFP mice ($n$ = 7) | ATm$^{VGluT2+}$-TeA/ECT-Arch mice ($n$ = 7) | Laser OFF | | | | |
| | | | Failed | | Mann-Whitney $U$ test (two-tailed) | $P$ = 0.3636 | $U$ = 15.5 |
| | | | Laser ON | | | | |
| | | | Failed | | Mann-Whitney $U$ test (two-tailed) | $P$ = 0.0023 | $U$ = 2 |
| Fig. 4i (right, Laser OFF vs Laser ON) | | ATm$^{VGluT2+}$-TeA/ECT-Arch mice ($n$ = 7) | Failed | | Wilcoxon matched-pairs signed rank test (two-tailed) | $P$ = 0.0313 | $W$ = -21 |
| | ATm$^{VGluT2+}$-TeA/ECT-eYFP mice ($n$ = 7) | | Failed | | Wilcoxon matched-pairs signed rank test (two-tailed) | $P$ = 0.1250 | $W$ = 13 |
| Fig. 4i (right, eYFP vs Arch) | ATm$^{VGluT2+}$-TeA/ECT-eYFP mice ($n$ = 7) | ATm$^{VGluT2+}$-TeA/ECT-Arch mice ($n$ = 7) | Laser OFF | | | | |
| | | | Failed | | Mann-Whitney $U$ test (two-tailed) | $P$ = 0.7308 | $U$ = 21 |
| | | | Laser ON | | | | |
| | | | Failed | | Mann-Whitney $U$ test (two-tailed) | $P$ = 0.0029 | $U$ = 2.5 |
| Fig. 4j (left, Laser OFF vs Laser ON) | | ATm$^{VGluT2+}$-TeA/ECT-Arch mice ($n$ = 7) | Failed | | Wilcoxon matched-pairs signed rank test (two-tailed) | $P$ = 0.0156 | $W$ = -28 |
| | ATm$^{VGluT2+}$-TeA/ECT-eYFP mice ($n$ = 4) | | Failed | | Wilcoxon matched-pairs signed rank test (two-tailed) | $P$ = 0.5000 | $W$ = -3 |
| Fig. 4j (left, eYFP vs Arch) | ATm$^{VGluT2+}$-TeA/ECT-eYFP mice ($n$ = 4) | ATm$^{VGluT2+}$-TeA/ECT-Arch mice ($n$ = 7) | Laser OFF | | | | |
| | | | Failed | | Mann-Whitney $U$ test (two-tailed) | $P$ = 0.0636 | $U$ = 4 |
| | | | Laser ON | | | | |
| | | | Failed | | Mann-Whitney $U$ test (two-tailed) | $P$ = 0.0030 | $U$ = 0 |
| Fig. 4j (right, Laser OFF vs Laser ON) | | ATm$^{VGluT2+}$-TeA/ECT-Arch mice ($n$ = 7) | Failed | | Wilcoxon matched-pairs signed rank test (two-tailed) | $P$ = 0.0313 | $W$ = -21 |
| | ATm$^{VGluT2+}$-TeA/ECT-eYFP mice ($n$ = 4) | | Passed | | Paired $t$ test (two-tailed) | $P$ = 0.0622 | $t_3$ = 2.905 |
| | | | Laser OFF | | | | |

| | | | | | | | |
|---|---|---|---|---|---|---|---|
| Fig. 4j (right, eYFP vs Arch) | ATm$^{VGluT2+}$-TeA/ECT-eYFP mice (*n* = 4) | ATm$^{VGluT2+}$-TeA/ECT-Arch mice (*n* = 7) | Failed | | Mann-Whitney *U* test (two-tailed) | $P$ = 0.3485 | $U$ = 8 |
| | | | Laser ON | | | | |
| | | | Failed | | Mann-Whitney *U* test (two-tailed) | $P$ = 0.0182 | $U$ = 2.5 |
| Fig. 5c (right, photoact. onset vs. photoact. offset) | | ATm$^{VGluT2+}$-ChR2 mice (*n* = 4) | Passed | | Paired *t* test (two-tailed) | $P$ = 0.0118 | $t_3$ = 5.511 |
| Fig. 5e (left, OF: No. of entries into center zone) | ATm$^{VGluT2+}$-eGFP mice (*n* = 6) | ATm$^{VGluT2+}$-ChR2 mice (*n* = 5) | Passed | Passed | Unpaired *t* test (two-tailed) | Pre-OFF: $P$ = 0.8317<br>ON: $P$ = 0.0002<br>Post-OFF: $P$ = 0.3161 | Pre-OFF: $t_9$ = 0.2188<br>ON: $t_9$ = 5.904<br>Post-OFF: $t_9$ = 1.062 |
| Fig. 5e (middle, OF: Time spent in center zone) | ATm$^{VGluT2+}$-eGFP mice (*n* = 6) | ATm$^{VGluT2+}$-ChR2 mice (*n* = 5) | Passed | Passed | Unpaired *t* test (two-tailed) | Pre-OFF: $P$ = 0.9017<br>ON: $P$ = 0.0172<br>Post-OFF: $P$ = 0.9660 | Pre-OFF: $t_9$ = 0.1271<br>ON: $t_9$ = 2.914<br>Post-OFF: $t_9$ = 0.04378 |
| Fig. 5e (right, OF: Average speed) | ATm$^{VGluT2+}$-eGFP mice (*n* = 6) | ATm$^{VGluT2+}$-ChR2 mice (*n* = 5) | Passed | Passed | Unpaired *t* test (two-tailed) | Pre-OFF: $P$ = 05333<br>ON: $P$ = 0.0041<br>Post-OFF: $P$ = 0.8067 | Pre-OFF: $t_9$ = 0.6477<br>ON: $t_9$ = 3.817<br>Post-OFF: $t_9$ = 0.2520 |
| Fig. 5g (left, EPM: No. of entries into OA) | ATm$^{VGluT2+}$-eGFP mice (*n* = 5) | ATm$^{VGluT2+}$-ChR2 mice (*n* = 5) | | | Pre-OFF | | |
| | | | Failed | | Mann-Whitney *U* test (two-tailed) | $P$ = 0.1746 | $U$ = 5.5 |
| | | | | | ON | | |
| | | | Passed | Passed | Unpaired *t* test (two-tailed) | $P$ = 0.0121 | $t_8$ = 3.225 |
| | | | | | Post-OFF | | |
| | | | | | Unpaired *t* test (two-tailed) | $P$ = 0.5149 | $t_8$ = 0.6814 |
| Fig. 5g (middle, EPM: Time spent in OA) | ATm$^{VGluT2+}$-eGFP mice (*n* = 5) | ATm$^{VGluT2+}$-ChR2 mice (*n* = 5) | Passed | Passed | Unpaired *t* test (two-tailed) | Pre-OFF: $P$ = 0.5116<br>ON: $P$ = 0.0003<br>Post-OFF: $P$ = 0.2765 | Pre-OFF: $t_8$ = 0.6868<br>ON: $t_8$ = 6.180<br>Post-OFF: $t_8$ = 1.168 |
| Fig. 5g (right, EPM: Average speed) | ATm$^{VGluT2+}$-eGFP mice (*n* = 5) | ATm$^{VGluT2+}$-ChR2 mice (*n* = 5) | | | Pre-OFF | | |
| | | | Passed | Passed | Unpaired *t* test (two-tailed) | $P$ = 0.2445 | $t_8$ = 1.256 |
| | | | | | ON | | |
| | | | Failed | | Mann-Whitney *U* test (two-tailed) | $P$ = 0.0079 | $U$ = 0 |
| | | | | | Post-OFF | | |
| | | | Passed | Passed | Unpaired *t* test (two-tailed) | $P$ = 0.6444 | $t_8$ = 0.4795 |
| Fig. 6d (Peak | ICx-ATm- | ICx-ATm- | | | Unpaired *t* | Pre (-2-0 s): $P$ = 0.2728<br>Noise (0-2 s): | Pre (-2-0 s): $t_4$ = 1.270<br>Noise (0-2 s): |

| | | | | | | | |
|---|---|---|---|---|---|---|---|
| speed: eYFP vs TeNT) | eYFP mice (*n* = 3) | TeNT mice (*n* = 3) | Passed | Passed | test (two-tailed) | [illegible] $P$ = 0.0106<br>Noise (2-10 s): $P$ = 0.0078 | [illegible] $t_4$ = 4.529<br>Noise (2-10 s): $t_4$ = 4.935 |
| Fig. 6e (Latency to escape: eYFP vs TeNT) | ICx-ATm-eYFP mice (n = 3 | ICx-ATm-TeNT mice (n = 3) | Passed | Passed | Unpaired t test (two-tailed) | $P$ = 0.0917 | $t_4$ = 2.209 |
| Fig. 6f (Time spent in shelter: eYFP vs TeNT) | ICx-ATm-eYFP mice (n = 3 | ICx-ATm-TeNT mice (n = 3) | Passed | Passed | Unpaired t test (two-tailed) | $P$ = 0.0411 | $t_4$ = 2.970 |

### Supplementary Figures

| Figure | *n* number | | Normality (Shapiro-Wilk test) | Equal variance test | Statistic method | *P* value | Statistics value |
|---|---|---|---|---|---|---|---|
| Supplementary Fig. 1c (right, UAN vs AN) | $ATm^{VGluT2+}$-GCaMP6m mice (*n* = 6) | | Failed | | Wilcoxon matched-pairs signed rank test (two-tailed) | $P$ = 0.0313 | $W$ = 21 |
| Supplementary Fig. 1c (right, UAN vs W) | $ATm^{VGluT2+}$-GCaMP6m mice (*n* = 6) | | Failed | | Mann-Whitney *U* test (two-tailed) | $P$ = 0.0152 | $U$ = 3 |
| Supplementary Fig. 1c (right, AN vs W) | $ATm^{VGluT2+}$-GCaMP6m mice (*n* = 6) | | Failed | | Mann-Whitney *U* test (two-tailed) | $P$ = 0.0931 | $U$ = 7 |
| Supplementary Fig. 1d (right, UAN vs AN) | $ATm^{VGluT2+}$-eGFP mice (*n* = 3) | | Failed | | Wilcoxon matched-pairs signed rank test (two-tailed) | $P$ > 0.9999 | $W$ = 0.0 |
| Supplementary Fig. 1d (right, UAN vs W) | $ATm^{VGluT2+}$-eGFP mice (*n* = 3) | | Failed | | Mann-Whitney *U* test (two-tailed) | $P$ = 0.7 | $U$ = 3 |
| Supplementary Fig. 1d (right, AN vs W) | $ATm^{VGluT2+}$-eGFP mice (*n* = 3) | | Failed | | Mann-Whitney *U* test (two-tailed) | $P$ = 0.7 | U = 3 |
| Supplementary Fig. 1e (right, NREM baseline vs AN) | MGBv-GCaMP6m mice (*n* = 5) | | Passed | | Paired *t* test (two-tailed) | $P$ = 0.6023 | $t_4$ = 0.5649 |
| Supplementary Fig. 1f (right, UAN vs AN) | MGBv-eGFP mice (*n* = 3) | | Passed | | Paired *t* test (two-tailed) | $P$ = 0.2501 | $t_2$ = 1.603 |
| Supplementary Fig. 1f (right, UAN vs W) | MGBv-eGFP mice (*n* = 3) | | Failed | | Mann-Whitney *U* test (two-tailed) | $P$ = 0.2000 | $U$ = 1 |
| Supplementary Fig. 1f (right, AN vs W) | MGBv-eGFP mice (*n* = 3) | | Failed | | Mann-Whitney *U* test (two-tailed) | $P$ > 0.9999 | $U$ = 4 |
| Supplementary Fig. 1g (right, NREM baseline vs AN) | MGBv-eGFP mice (*n* = 3) | | Passed | | Paired *t* test (two-tailed) | $P$ = 0.2746 | $t_2$ = 1.491 |
| Supplementary Fig. 2c (right, NREM) | MGBv-eGFP mice (*n* = 4) | MGBv-ChR2 mice (*n* = 6) | Passed | Passed | Unpaired *t* test (two-tailed) | $P$ = 0.5877 | $t_8$ = 0.5648 |
| Supplementary Fig. 2c (right, REM) | MGBv-eGFP mice (*n* = 4) | MGBv-ChR2 mice (*n* = 6) | Passed | Passed | Unpaired *t* test (two-tailed) | $P$ = 0.0791 | $t_8$ = 2.011 |
| Supplementary Fig. 2c (right, Wake) | MGBv-eGFP mice (*n* = 4) | MGBv-ChR2 mice (*n* = 6) | Passed | Passed | Unpaired *t* test (two-tailed) | $P$ = 0.6906 | $t_8$ = 0.4129 |

| | | | | | | | |
|---|---|---|---|---|---|---|---|
| Supplementary Fig. 2g (right, NREM) | dLGN-eGFP mice (*n* = 5) | dLGN-ChR2 mice (*n* = 5) | Passed | Passed | Unpaired *t* test (two-tailed) | $P$ = 0.3837 | $t_8$ = 0.9215 |
| Supplementary Fig. 2g (right, REM) | dLGN-eGFP mice (*n* = 5) | dLGN-ChR2 mice (*n* = 5) | Passed | Passed | Unpaired *t* test (two-tailed) | $P$ = 0.0582 | $t_8$ = 2.209 |
| Supplementary Fig. 2g (right, Wake) | dLGN-eGFP mice (*n* = 5) | dLGN-ChR2 mice (*n* = 5) | Passed | Passed | Unpaired *t* test (two-tailed) | $P$ = 0.9328 | $t_8$ = 0.08702 |
| Supplementary Fig. 2h (right, NREM) | dLGN-eGFP mice (*n* = 4) | dLGN-ChR2 mice (*n* = 5) | Passed | Passed | Unpaired *t* test (two-tailed) | $P$ = 0.7824 | $t_7$ = 0.2870 |
| Supplementary Fig. 2h (right, REM) | dLGN-eGFP mice (*n* = 4) | dLGN-ChR2 mice (*n* = 5) | Passed | Passed | Unpaired *t* test (two-tailed) | $P$ = 0.2528 | $t_7$ = 1.246 |
| Supplementary Fig. 2h (right, Wake) | dLGN-eGFP mice (*n* = 4) | dLGN-ChR2 mice (*n* = 5) | Passed | Passed | Unpaired *t* test (two-tailed) | $P$ = 03372 | $t_7$ = 1.030 |
| Supplementary Fig. 3c (right, NREM) | $ICx^{VGluT2+}$-ATm-eGFP mice (*n* = 4) | $ICx^{VGluT2+}$-ATm-ChR2 mice (*n* = 5) | Passed | Passed | Unpaired *t* test (two-tailed) | $P$ < 0.0001 | $t_7$ = 11.32 |
| Supplementary Fig. 3c (right, REM) | $ICx^{VGluT2+}$-ATm-eGFP mice (*n* = 4) | $ICx^{VGluT2+}$-ATm-ChR2 mice (*n* = 5) | Passed | Passed | Unpaired *t* test (two-tailed) | $P$ = 0.6781 | $t_7$ = 0.4329 |
| Supplementary Fig. 3c (right, Wake) | $ICx^{VGluT2+}$-ATm-eGFP mice (*n* = 4) | $ICx^{VGluT2+}$-ATm-ChR2 mice (*n* = 5) | Failed | | Mann-Whitney *U* test (two-tailed) | $P$ = 0.0159 | $U$ = 0 |
| Supplementary Fig. 3d (right, NREM) | $SO/dSC^{VGluT2+}$-ATm-eGFP mice (*n* = 6) | $SO/dSC^{VGluT2+}$-ATm-ChR2 mice (*n* = 4) | Passed | Passed | Unpaired *t* test (two-tailed) | $P$ < 0.0001 | $t_8$ = 18.96 |
| Supplementary Fig. 3d (right, REM) | $SO/dSC^{VGluT2+}$-ATm-eGFP mice (*n* = 6) | $SO/dSC^{VGluT2+}$-ATm-ChR2 mice (*n* = 4) | Passed | Passed | Unpaired *t* test (two-tailed) | $P$ = 0.8829 | $t_8$ = 0.1521 |
| Supplementary Fig. 3d (right, Wake) | $SO/dSC^{VGluT2+}$-ATm-eGFP mice (*n* = 6) | $SO/dSC^{VGluT2+}$-ATm-ChR2 mice (*n* = 4) | Passed | Passed | Unpaired *t* test (two-tailed) | $P$ < 0.0001 | $t_8$ = 17.59 |
| Supplementary Fig. 4e (left, 0-5 s) | $ATm^{VGluT2+}$-TeA/ECT-eYFP mice (*n* = 5) | $ATm^{VGluT2+}$-TeA/ECT-ChR2 mice (*n* = 7) | Failed | | Mann-Whitney *U* test (two-tailed) | $P$ = 0.029 | $U$ = 4.5 |
| Supplementary Fig. 4e (left, 5-10 s) | $ATm^{VGluT2+}$-TeA/ECT-eYFP mice (*n* = 5) | $ATm^{VGluT2+}$-TeA/ECT-ChR2 mice (*n* = 7) | Failed | | Mann-Whitney *U* test (two-tailed) | $P$ = 0.0025 | $U$ = 0 |
| Supplementary Fig. 4e (left, 10-20 s) | $ATm^{VGluT2+}$-TeA/ECT-eYFP mice (*n* = 5) | $ATm^{VGluT2+}$-TeA/ECT-ChR2 mice (*n* = 7) | Passed | Passed | Unpaired *t* test (two-tailed) | $P$ = 0.1516 | $t_{10}$ = 1.553 |
| Supplementary Fig. 4e (left, >20 s) | $ATm^{VGluT2+}$-TeA/ECT-eYFP mice (*n* = 5) | $ATm^{VGluT2+}$-TeA/ECT-ChR2 mice (*n* = 7) | Passed | Passed | Unpaired *t* test (two-tailed) | $P$ = 0.0001 | $t_{10}$ = 5.915 |
| Supplementary Fig. 4e (right, 0-5 s) | $ATm^{VGluT2+}$-TeA/ECT-eYFP mice (*n* = 5) | $ATm^{VGluT2+}$-TeA/ECT-ChR2 mice (*n* = 7) | Failed | | Mann-Whitney *U* test (two-tailed) | $P$ = 0.0316 | $U$ = 4 |
| Supplementary Fig. 4e (right, 5-10 s) | $ATm^{VGluT2+}$-TeA/ECT-eYFP mice (*n* = 5) | $ATm^{VGluT2+}$-TeA/ECT-ChR2 mice (*n* = 7) | Failed | | Mann-Whitney *U* test (two-tailed) | $P$ = 0.0114 | $U$ = 2.5 |
| Supplementary Fig. 4e (right, 10-20 s) | $ATm^{VGluT2+}$-TeA/ECT-eYFP mice (*n* = 5) | $ATm^{VGluT2+}$-TeA/ECT-ChR2 mice (*n* = 7) | Failed | | Mann-Whitney *U* test (two-tailed) | $P$ = 0.0657 | $U$ = 6 |

| | | | | | | | |
|---|---|---|---|---|---|---|---|
| Supplementary Fig. 4e (right, >20 s) | ATm$^{VGluT2+}$-TeA/ECT-eYFP mice ($n$ = 5) | ATm$^{VGluT2+}$-TeA/ECT-ChR2 mice ($n$ = 7) | Failed | | Mann-Whitney $U$ test (two-tailed) | $P$ = 0.0025 | $U$ = 0 |
| Supplementary Fig. 5b (left, Laser OFF vs Laser ON) | | ATm$^{VGluT2+}$-VMH-Arch mice ($n$ = 7) | Failed | | Wilcoxon matched-pairs signed rank test (two-tailed) | $P$ = 0.0313 | $W$ = -21 |
| | ATm$^{VGluT2+}$-VMH-eGFP mice ($n$ = 5) | | Failed | | Wilcoxon matched-pairs signed rank test (two-tailed) | $P$ = 0.5000 | $W$ = 4 |
| Supplementary Fig. 5b (left, eGFP vs Arch) | ATm$^{VGluT2+}$-VMH-eGFP mice ($n$ = 5) | ATm$^{VGluT2+}$-VMH-Arch mice ($n$ = 7) | Laser OFF | | | | |
| | | | Failed | | Mann-Whitney $U$ test (two-tailed) | $P$ > 0.9999 | $U$ = 17 |
| | | | Laser ON | | | | |
| | | | Failed | | Mann-Whitney $U$ test (two-tailed) | $P$ = 0.0253 | $U$ = 4 |
| Supplementary Fig. 5b (right, Laser OFF vs Laser ON) | | ATm$^{VGluT2+}$-VMH-Arch mice ($n$ = 7) | Failed | | Wilcoxon matched-pairs signed rank test (two-tailed) | $P$ = 0.0313 | $W$ = -21 |
| | ATm$^{VGluT2+}$-VMH-eGFP mice ($n$ = 5) | | Passed | | Paired $t$ test (two-tailed) | $P$ = 0.3739 | $t_4$ = 1 |
| Supplementary Fig. 5b (right, eGFP vs Arch) | ATm$^{VGluT2+}$-VMH-eGFP mice ($n$ = 5) | ATm$^{VGluT2+}$-VMH-Arch mice ($n$ = 7) | Laser OFF | | | | |
| | | | Passed | Passed | Unpaired $t$ test (two-tailed) | $P$ = 0.8878 | $t_{10}$ = 0.1447 |
| | | | Laser ON | | | | |
| | | | Failed | | Mann-Whitney $U$ test (two-tailed) | $P$ = 0.0202 | $U$ = 5 |
| Supplementary Fig. 5c (left, Laser OFF vs Laser ON) | | ATm$^{VGluT2+}$-VMH-Arch mice ($n$ = 7) | Failed | | Wilcoxon matched-pairs signed rank test (two-tailed) | $P$ = 0.0625 | $W$ = -15 |
| | ATm$^{VGluT2+}$-VMH-eGFP mice ($n$ = 5) | | Failed | | Wilcoxon matched-pairs signed rank test (two-tailed) | $P$ > 0.9999 | $W$ = 2 |
| Supplementary Fig. 5c (left, eGFP vs Arch) | ATm$^{VGluT2+}$-VMH-eGFP mice ($n$ = 5) | ATm$^{VGluT2+}$-VMH-Arch mice ($n$ = 7) | Laser OFF | | | | |
| | | | Failed | | Mann-Whitney $U$ test (two-tailed) | $P$ = 0.5581 | $U$ = 13 |
| | | | Laser ON | | | | |
| | | | Failed | | Mann-Whitney $U$ test (two-tailed) | $P$ = 0.1111 | $U$ = 7.5 |
| Supplementary Fig. 5c (right, Laser OFF vs Laser ON) | | ATm$^{VGluT2+}$-VMH-Arch mice ($n$ = 7) | Failed | | Wilcoxon matched-pairs signed rank test (two-tailed) | $P$ = 0.3125 | $W$ = -10 |
| | ATm$^{VGluT2+}$-VMH-eGFP mice ($n$ = 5) | | Passed | | Paired $t$ test (two-tailed) | $P$ = 0.0533 | $t_4$ = 2.714 |
| Supplementary Fig. 5c (right, eGFP vs Arch) | ATm$^{VGluT2+}$-VMH-eGFP mice ($n$ = 5) | ATm$^{VGluT2+}$-VMH-Arch mice ($n$ = [illegible] | Laser OFF | | | | |
| | | | Passed | Passed | Unpaired $t$ test (two-tailed) | $P$ = 0.2057 | $t_{10}$ = 1.354 |
| | | | Laser ON | | | | |

| | | | | | | | |
|---|---|---|---|---|---|---|---|
| | | 7) | Failed | | Mann-Whitney *U* test (two-tailed) | $P$ = 0.2058 | $U$ = 9 |
| Supplementary Fig. 5d (left, Laser OFF vs Laser ON) | | ATm$^{VGluT2+}$-TS-Arch mice ($n$ = 6) | Passed | | Paired *t* test (two-tailed) | $P$ = 0.1019 | $t_5$ = 2.000 |
| | ATm$^{VGluT2+}$-TS-eGFP mice ($n$ = 5) | | Failed | | Wilcoxon matched-pairs signed rank test (two-tailed) | $P >$ 0.9999 | $W$ = 1 |
| Supplementary Fig. 5d (left, eGFP vs Arch) | ATm$^{VGluT2+}$-TS-eGFP mice ($n$ = 5) | ATm$^{VGluT2+}$-TS-Arch mice ($n$ = 6) | Laser OFF | | | | |
| | | | Failed | | Mann-Whitney *U* test (two-tailed) | $P$ = 0.4589 | $U$ = 10 |
| | | | Laser ON | | | | |
| | | | Failed | | Mann-Whitney *U* test (two-tailed) | $P$ = 0.4589 | $U$ = 10 |
| Supplementary Fig. 5d (right, Laser OFF vs Laser ON) | | ATm$^{VGluT2+}$-TS-Arch mice ($n$ = 6) | Passed | | Paired *t* test (two-tailed) | $P$ = 0.2242 | $t_5$ = 1.387 |
| | ATm$^{VGluT2+}$-TS-eGFP mice ($n$ = 5) | | Passed | | Paired *t* test (two-tailed) | $P$ = 0.6213 | $t_4$ = 0.5345 |
| Supplementary Fig. 5d (right, eGFP vs Arch) | ATm$^{VGluT2+}$-TS-eGFP mice ($n$ = 5) | ATm$^{VGluT2+}$-TS-Arch mice ($n$ = 6) | Laser OFF | | | | |
| | | | Passed | Passed | Unpaired *t* test (two-tailed) | $P$ = 0.1301 | $t_9$ = 1.666 |
| | | | Laser ON | | | | |
| | | | Passed | Passed | Unpaired *t* test (two-tailed) | $P$ = 0.9212 | $t_9$ = 0.1018 |
| Supplementary Fig. 5e (left, Laser OFF vs Laser ON) | | ATm$^{VGluT2+}$-TS-Arch mice ($n$ = 6) | Failed | | Wilcoxon matched-pairs signed rank test (two-tailed) | $P$ = 0.0313 | $W$ = -21 |
| | ATm$^{VGluT2+}$-TS-eGFP mice ($n$ = 5) | | Passed | | Paired *t* test (two-tailed) | $P$ > 0.9999 | $t_4$ = 3.511e-016 |
| Supplementary Fig. 5e (left, eGFP vs Arch) | ATm$^{VGluT2+}$-TS-eGFP mice ($n$ = 5) | ATm$^{VGluT2+}$-TS-Arch mice ($n$ = 6) | Laser OFF | | | | |
| | | | Failed | | Mann-Whitney *U* test (two-tailed) | $P$ = 0.3636 | $U$ = 9 |
| | | | Laser ON | | | | |
| | | | Failed | | Mann-Whitney *U* test (two-tailed) | $P$ = 0.0433 | $U$ = 3 |
| Supplementary Fig. 5e (right, Laser OFF vs Laser ON) | | ATm$^{VGluT2+}$-TS-Arch mice ($n$ = 6) | Passed | | Paired *t* test (two-tailed) | $P$ = 0.4017 | $t_5$ = 0.9159 |
| | ATm$^{VGluT2+}$-TS-eGFP mice ($n$ = 5) | | Passed | | Paired *t* test (two-tailed) | $P$ = 0.4766 | $t_4$ = 0.7845 |
| | | | Laser OFF | | | | |

| | | | | | | | |
|---|---|---|---|---|---|---|---|
| Supplementary Fig. 5e (right, eGFP vs Arch) | $ATm^{VGluT2+}$-TS-eGFP mice (*n* = 5) | $ATm^{VGluT2+}$-TS-Arch mice (*n* = 6) | Passed | Passed | Unpaired *t* test (two-tailed) | *P* = 0.8243 | $t_9$ =0.2285 |
| | | | Laser ON | | | | |
| | | | Passed | Passed | Unpaired *t* test (two-tailed) | *P* = 0.0594 | $t_9$ =2.157 |
| Supplementary Fig. 6b (left, RTPA: No. of entries into Stim zone) | $ATm^{VGluT2+}$-eGFP mice (*n* = 6) | $ATm^{VGluT2+}$-ChR2 mice (*n* = 5) | Passed | Passed | Unpaired *t* test (two-tailed) | Pre-OFF: *P* = 0.7187<br>ON: *P* = 0.0027<br>Post-OFF: *P* = 0.3661 | Pre-OFF: $t_9$ = 0.3717<br>ON: $t_9$ = 4.087<br>Post-OFF: $t_9$ = 0.9517 |
| Supplementary Fig. 6b (middle, RTPA: Time spent in Stim zone) | $ATm^{VGluT2+}$-eGFP mice (*n* = 6) | $ATm^{VGluT2+}$-ChR2 mice (*n* = 5) | Passed | Passed | Unpaired *t* test (two-tailed) | Pre-OFF: *P* = 0.2036<br>ON: *P* < 0.0001<br>Post-OFF: *P* = 0.3797 | Pre-OFF: $t_9$ = 1.371<br>ON: $t_9$ = 7.799<br>Post-OFF: $t_9$ = 0.9238 |
| Supplementary Fig. 6b (right, RTPA: Average speed) | $ATm^{VGluT2+}$-eGFP mice (*n* = 6) | $ATm^{VGluT2+}$-ChR2 mice (*n* = 5) | Passed | Passed | Unpaired *t* test (two-tailed) | Pre-OFF: *P* = 0.5991<br>ON: *P* = 0.0007<br>Post-OFF: *P* = 0.2995 | Pre-OFF: $t_9$ = 0.5449<br>ON: $t_9$ = 5.051<br>Post-OFF: $t_9$ = 1.101 |
